# Functional testing of PI3K inhibitors stratifies responders to idelalisib and identifies treatment vulnerabilities in idelalisib-refractory/intolerant chronic lymphocytic leukemia

**DOI:** 10.1101/2022.04.14.488428

**Authors:** Yanping Yin, Paschalis Athanasiadis, Linda Karlsen, Aleksandra Urban, Ishwarya Murali, Stacey M. Fernandes, Alberto J. Arribas, Abdul K. Hilli, Kjetil Taskén, Francesco Bertoni, Anthony R. Mato, Emmanuel Normant, Jennifer R. Brown, Geir E. Tjønnfjord, Tero Aittokallio, Sigrid S. Skånland

**Author notes:** **Correspondence:** Sigrid S. Skånland, Department of Cancer Immunology, Institute for Cancer Research, Oslo University Hospital, P.O. Box 4951 Nydalen, N-0424, Norway.

## Abstract

**Purpose:** Phosphatidylinositol 3-kinase inhibitors (PI3Ki) are approved for relapsed chronic lymphocytic leukemia (CLL). While patients may show an initial response, development of treatment intolerance or resistance remains a clinically challenging. Prediction of individual treatment responses based on clinically actionable biomarkers is needed to overcome these challenges. Here, we investigated whether *ex vivo* functional responses to targeted therapies can stratify responders to idelalisib and guide precision medicine in CLL.

**Experimental design:** CLL cells from treatment naïve, idelalisib-responding, and idelalisib-refractory/intolerant patients (n=33 in total) were profiled against ten PI3Ki and the Bcl-2 antagonist venetoclax. Cell signaling and immune phenotypes were analyzed by flow cytometry. Cell viability was monitored by detection of cleaved caspase-3 and the CellTiter-Glo assay.

**Results:** Among the ten PI3Ki studied, pan-PI3Ki were most effective at inhibiting PI3K signaling and cell viability, and they showed activity also in CLL cells from idelalisib-refractory/intolerant patients. The pan-PI3Ki copanlisib, but not the p110δ inhibitor idelalisib, inhibited PI3K signaling in CD4^+^ and CD8^+^ T cells in addition to CD19^+^ B cells, while it did not significantly affect T cell numbers. Combination treatment with a PI3Ki and venetoclax resulted in synergistic induction of apoptosis. Based on *ex vivo* drug sensitivity testing, a relapsed CLL patient was treated with idelalisib plus venetoclax, and the patient achieved a partial response. A more systematic analysis revealed that CLL cells from patients with a long-term response to idelalisib showed significantly higher drug sensitivities to 73 drug combinations at baseline compared to short-term responders.

**Conclusions:** Our findings suggest novel treatment vulnerabilities in idelalisib-refractory/intolerant CLL, and demonstrate that *ex vivo* functional profiling may guide precision medicine and predict treatment responses of individual CLL patients.

**TRANSLATIONAL RELEVANCE:** The phosphatidylinositol 3-kinase inhibitors (PI3Ki) idelalisib and duvelisib are approved for relapsed chronic lymphocytic leukemia (CLL), but their use has been limited by severe toxicity and acquired resistance. Identification of biomarkers that predict individual treatment responses, as well as alternative treatment vulnerabilities in PI3Ki refractory/intolerant patients, is needed to optimally tailor CLL therapy. We performed functional analyses of CLL cells from treatment naïve, idelalisib-responding and idelalisib-refractory/intolerant patients to identify clinically actionable biomarkers. We show that CLL cells from idelalisib-refractory/intolerant patients remain sensitive to pan-PI3Ki and PI3Ki plus venetoclax combinations. *Ex vivo* drug sensitivity testing was used to guide treatment of a relapsed CLL patient who obtained a partial response after idelalisib plus venetoclax therapy. A systematic analysis of drug sensitivities to 73 drug combinations stratified responders to idelalisib using baseline samples from short-term and long-term responders to idelalisib. Our study demonstrates the power of functional precision medicine in relapsed CLL.

## INTRODUCTION

Excessive and autonomous B cell receptor (BCR) signaling is an important pathogenic mechanism in CLL,^1^ where continuous signaling leads to activation of downstream signaling molecules such as phosphatidylinositol 3-kinase (PI3K).^2^ Among the four classes of PI3Ks, only the class I isoforms are clearly implicated in cancer.^3^ The catalytic subunit of class I PI3K exists as different isoforms, p110α, β, γ and δ. The α and β isoforms are ubiquitously expressed, while γ and δ are primarily expressed in leukocytes.^4^ P110δ is highly expressed in CLL cells, which makes it a promising target for CLL treatment.^5^ So far, two PI3K inhibitors (PI3Ki), idelalisib and duvelisib, have been approved for treatment of CLL, and several next-generation inhibitors are in development.^6^

Even though targeted therapies have revolutionized the management of CLL, treatment intolerance and resistance remain clinically challenging.^7;8^ To date, no mutations in PI3Ks are known that could explain the resistance mechanism to PI3Ki.^9;10^ Rather, reports suggest that resistance to PI3Ki may be mediated by upregulation of p110 isoforms other than p110δ^11-13^ This supports the use of a pan-p110 inhibitor (here referred to as pan-PI3Ki) such as copanlisib, which is approved for indolent lymphoma. Upregulation of alternative signaling pathways may also explain the development of PI3Ki resistance in some cases. Activation of STAT3 or STAT5 by interleukin-6 or ectopic activation of the ERBB signaling were shown to underlie resistance to various PI3Ki in lymphoma,^14;15^ while activating mutations in MAPK pathway genes and upregulation of IGF1R are reported resistance mechanisms in CLL.^16;17^ These resistance mechanisms provide a rationale for combination therapies, and we recently showed that MEK inhibitors are effective in idelalisib-resistant CLL.^18^

Despite the initially encouraging results for PI3Ki in CLL, a more widespread use of approved PI3Ki has been limited by unexpected autoimmune toxicities.^5;7^ Current strategies to allow for optimized use of the drug class include exploring next-generation PI3Ki and alternative dosing schedules.^19^ Prospective clinical trials have demonstrated that patients who experience intolerance or resistance to a targeted therapy may still benefit from another drug within the same drug class.^20-22^ However, there is currently a limited knowledge of how the overall sensitivity to PI3Ki evolves in response to idelalisib treatment, and whether idelalisib-refractory/intolerant CLL cells remain sensitive to other PI3Ki. *Ex vivo* drug testing in patient cells has recently demonstrated clinical utility in guiding functional precision medicine for CLL and other hematological malignancies.^23-25^ When combined with other functional assays, such an integrated approach can lead to actionable biomarkers that predict patient treatment responses and identify unexpected treatment vulnerabilities such as novel combinations.^26^

Here, we combined *ex vivo* profiling of drug sensitivity and cell signaling in response to ten different PI3Ki (buparlisib, compound 7n, copanlisib, duvelisib, idelalisib, nemiralisib, pictilisib, pilaralisib, umbralisib, ZSTK474) in CLL, both alone and in combination with the B-cell lymphoma-2 (Bcl-2) antagonist venetoclax. We found that pan-PI3Ki were most effective in inhibiting cell signaling and viability of CLL cells. Furthermore, pan-PI3Ki single agents and PI3Ki + venetoclax combinations showed activity both in idelalisib-resistant lymphoma cell lines and in CLL cells from idelalisib-refractory/intolerant patients. To demonstrate its clinical feasibility, *ex vivo* drug testing was used to guide treatment with idelalisib + venetoclax of a relapsed CLL patient who had progressive disease after several lines of therapy. A more systematic analysis of longitudinal samples from idelalisib-treated patients demonstrated significantly higher *ex vivo* combination sensitivities at baseline in patients who obtained a long-term response to idelalisib.

Our findings indicate PI3Ki drug class activity in idelalisib-refractory/intolerant CLL and suggest that functional precision medicine based on *ex vivo* testing of drug sensitivity and cell signaling provides an exciting opportunity to predict treatment responses and to identify potential treatment options, both monotherapies and combinations. These results warrant further testing in larger cohorts and in clinical trials.

## MATERIALS AND METHODS

### Patient material and ethical considerations

Buffy coats from age-matched, anonymized healthy blood donors were received from the Department of Immunology and Transfusion Medicine, Oslo University Hospital, Norway. Blood samples from CLL patients were received from the Department of Haematology, Oslo University Hospital, Norway; the Department of Medicine, Diakonhjemmet Hospital, Norway; Dana-Farber Cancer Institute (DFCI), MA, USA; and TG Therapeutics, New York, NY, USA (NCT02742090). All participants signed a written informed consent prior to sample collection. The study was approved by the Regional Committee for Medical and Health Research Ethics of South-East Norway (2016/947 and 28507). The DFCI tissue bank protocol was approved by the Dana-Farber Harvard Cancer Center Institutional Review Board. The NCT02742090 study was done in compliance with good clinical practice and local and national regulatory guidelines. An institutional review board at each site approved the protocol before any patients were enrolled. Research on blood samples was carried out in agreement with the Declaration of Helsinki.

### Reagents and antibodies

Venetoclax, buparlisib, copanlisib, compound 7n, duvelisib, idelalisib, nemiralisib, pictilisib, pilaralisib, umbralisib and ZSTK474 were obtained from Selleckchem (Houston, TX, USA). All compounds and combinations used in this study are listed in **Supplementary Table 1** and **Supplementary Table 2**, respectively. Antibodies against AKT (pS473) (D9E), cleaved caspase-3 (Asp175) (D3PE), NF-κB p65 (pS536) (93H1), p38 MAPK (pT180/Y182) (28B10), p44/42 MAPK (pT202/Y204) (E10), S6-ribosomal protein (pS235/S236) (D57.2.2E) and SYK (pY525/Y526) (C87C1) were from Cell Signaling Technologies (Leiden, The Netherlands). Antibodies against Btk (pY223)/Itk (pY180) (N35-86), Btk (pY551) & p-Itk (pY511) (24a/BTK (Y551), IgG1 Kappa (MOPC-21), MEK1 (pS298) (J114-64), mTOR (pS2448) (O21-404), PLCγ2 (pY759) (K86-689.37), STAT3 (pS727) (49/p-Stat3), ZAP70/Syk (pY319/Y352) (17A/P-ZAP70) were from BD Biosciences (San Jose, CA, USA). These antibodies were conjugated to Alexa Fluor 647. PerCP-Cy5.5 conjugated mouse anti-human CD19 antibody (HIB19) was from eBioscience (San Diego, CA, USA). PE-Cy7 conjugated mouse anti-human CD3 antibody (UCHT1) was from BD Biosciences. Goat F(ab’)2 anti-human IgM was from Southern Biotech (Birmingham, AL, USA). BD Phosflow Fix Buffer I and Perm Buffer III were from BD Biosciences. Alexa Fluor 488, Pacific Blue and Pacific Orange Succinimidyl Esters were from Thermo Fisher Scientific (Waltham, MA, USA).

### Kinome scan

The activities of compound 7n, copanlisib, nemiralisib, and pilaralisib were profiled at 1 μM across a panel of 468 human kinases, including atypical, mutant, lipid, and pathogen kinases, using the DiscoveRx competition binding assay (Eurofins Scientific, Luxembourg). The measured percent of inhibition data are shown in **Supplementary Table 3**. Similar kinase activity data for buparlisib, duvelisib, idelalisib, pictilisib, umbralisib and ZSTK474, profiled following the same kinome scan protocol (https://www.discoverx.com/services/drug-discovery-development-services/kinase-profiling/kinomescan), but with a panel of 442 or 468 kinases, were available from previous studies.^27;28^ The activity data were visualized using the web-based tool TREE*spot* (Eurofins Scientific).

### Isolation of lymphocytes and peripheral blood mononuclear cells (PBMCs)

Isolation of CD19^+^ B cells from buffy coats from healthy blood donors and PBMCs from CLL patient samples was performed as previously described.^29^ Isolated cells were cryopreserved in liquid nitrogen.^30^ See **Supplementary Table 4** for patient characteristics.

### Phospho flow with fluorescent cell barcoding

Experiments were performed as previously described.^31;32^ For experiments shown in **Figure 2b-d** and **Figure 4a-c**, PBMCs from CLL153, CLL159, CLL160 and CLL216 (**Supplementary Table 4**) were treated with the indicated compound and concentration for 30 min, followed by 5 min stimulation with anti-IgM to activate the BCR. For experiments shown in **Figure 2e-j**, PBMCs from CLL002D, CLL216 and CLL248 (**Supplementary Table 4**) were simultaneously co-cultured with irradiated (50 Gy/125 Gy/125 Gy, respectively) CD40L^+^, BAFF^+^ and APRIL^+^ 3T3 fibroblasts (ratio 1:1:1) to prevent induction of spontaneous apoptosis,^33^ and treated with the indicated compound and concentration for 24h. The treated CLL cells were then fixed, barcoded and permeabilized as previously described.^31^ The samples were stained with the indicated antibodies and analyzed with a BD LSR Fortessa cytometer (BD Biosciences) equipped with 488 nm, 561 nm, 640 nm and 407 nm lasers. The data were analyzed in Cytobank (https://cellmass.cytobank.org/cytobank/) as previously described.^32^

### Immune phenotyping

PBMCs from each CLL donor were fixed, barcoded,^31^ and stained for 30 min with the antibody panel. The following antibodies were used: CD3-BUV395, CD4-BUV563, HLA-DR-BUV615, CD16-BUV737, CXCR5 (CD185)-BUV805, CCR7 (CD197)-BV605, CCR6 (CD196)-BV711, CD56-BV750, CD127-BV786, PD-1 (CD279)-BB700, CD14-PE, CD25-PE-CF594, CCR3 (CD183)-PE-Cy5, CD45RA-PE-Cy7, CD69-APC-R700, CD8/CD19-APC-Cy7 (BD Biosciences). Experiments were analyzed with a BD FACSymphony A5 cytometer (BD Biosciences) and further processed in Cytobank (https://cellmass.cytobank.org/cytobank/). FlowSOM clustering algorithm was applied to identify cell populations, which were validated by manual gating. The UMAP dimensionality reduction algorithm was used to visualize the data.

### Cell lines

Idelalisib-resistant lines from VL51 and KARPAS1718 *in vitro* models were developed by prolonged exposure to the IC90 concentration of idelalisib, as previously described.^15^

### CellTiter-Glo luminescent cell viability assay

Dose-response experiments were performed as previously described.^18;33^ Compounds (**Supplementary Table 1**) were pre-printed in 384-well cell culture microplates using the Echo 550 liquid handler (Labcyte Inc., San Jose, CA, USA). Each compound was tested at five different concentrations in ten-fold increments ranging from 1 nM - 10000 nM (0.1 nM - 1000 nM for copanlisib). Combinations were designed using a fixed molar concentration series identical to those used for single agents. PBMCs from CLL patient samples were co-cultured with irradiated (50 Gy/125 Gy/125 Gy, respectively) CD40L^+^, BAFF^+^ and APRIL^+^ 3T3 fibroblasts (ratio 1:1:1) for 24 h prior to initiation of the experiment to mimic the tumor microenvironment and to prevent spontaneous apoptosis. The CLL cells were then separated from the adherent fibroblast layer by carefully re-suspending the culturing medium and transferring it to a separate tube. Experiments on “JB” samples (**Supplementary Table 4**) were performed using a slightly modified protocol, in which non-irradiated fibroblasts were removed from the CLL cells by positive selection with a PE-conjugated anti-CD47 antibody and anti-PE microbeads, as previously described.^18^ Single-cell suspension (10000 cells/well in 25 μl) was distributed to each well of the 384-well assay plate using the CERTUS Flex liquid dispenser (Fritz Gyger, Thun, Switzerland). Dose-response experiments on cell lines were performed on freshly thawed cells to reduce variation between experiments introduced by culturing conditions. The cells were incubated with the compounds at 37°C for 72h. Cell viability was measured using the CellTiter-Glo luminescent assay (Promega, Madison, WI, USA) according to the manufacturer’s instructions. Luminescence was recorded with an EnVision 2102 Multilabel Reader (PerkinElmer, Waltham, MA, USA). The response readout was normalized to the negative (0.1% DMSO) and positive (100 µM benzethonium chloride) controls. The measured dose–response data were processed with the KNIME software (KNIME AG, Zurich, Switzerland).

### Data analyses

Measured data from phospho flow experiments and drug sensitivity screens were processed in GraphPad Prism 8 (San Diego, CA, USA). Applied statistical tests are indicated in the figure legends. The normality of the data distribution was tested using the Kolmogorov-Smirnov test in GraphPad Prism 8. To quantify the compound responses, a modified drug sensitivity score (DSS) was calculated for each sample and compound separately.^34^ Area under the dose-response curve was calculated using an activity window from 100% to 10%, and a dose-window from the minimum concentration tested to the concentration where the viability reached 10%. DSS_3_ metric was used, without the division by the logarithm of the upper asymptote of the logistic curve. Higher levels of DSS indicate higher sensitivity to the compound. The DECREASE tool was used to predict the full dose-response matrices based on the diagonal (dose ratio 1:1) combination experiments based on normalized phospho flow data.^33;35^ Combination synergy was scored using the Bliss model using the SynergyFinder web-tool.^36;37^ Figure 5a was created with BioRender.com.

## RESULTS

### Target specificity and activity profiles of PI3K inhibitors

The target activity of ten PI3Ki was profiled at 1 μM over a panel of up to 468 human kinases using the kinome scan assay (**Figure 1**), and includes earlier reported data.^27;28^ The inhibitors showed expected differences in their p110 isoform specificity and off-target profiles (**Figure 1** and **Supplementary Table 3**). Among the PI3Ki, compound 7n, nemiralisib and umbralisib showed p110δ isoform-specific activity with relatively few off-target activities, while copanlisib and ZSTK474 showed broad activity against all four p110 isoforms and several off-targets (**Figure 1** and **Figure 2a**).

**Figure 1.**
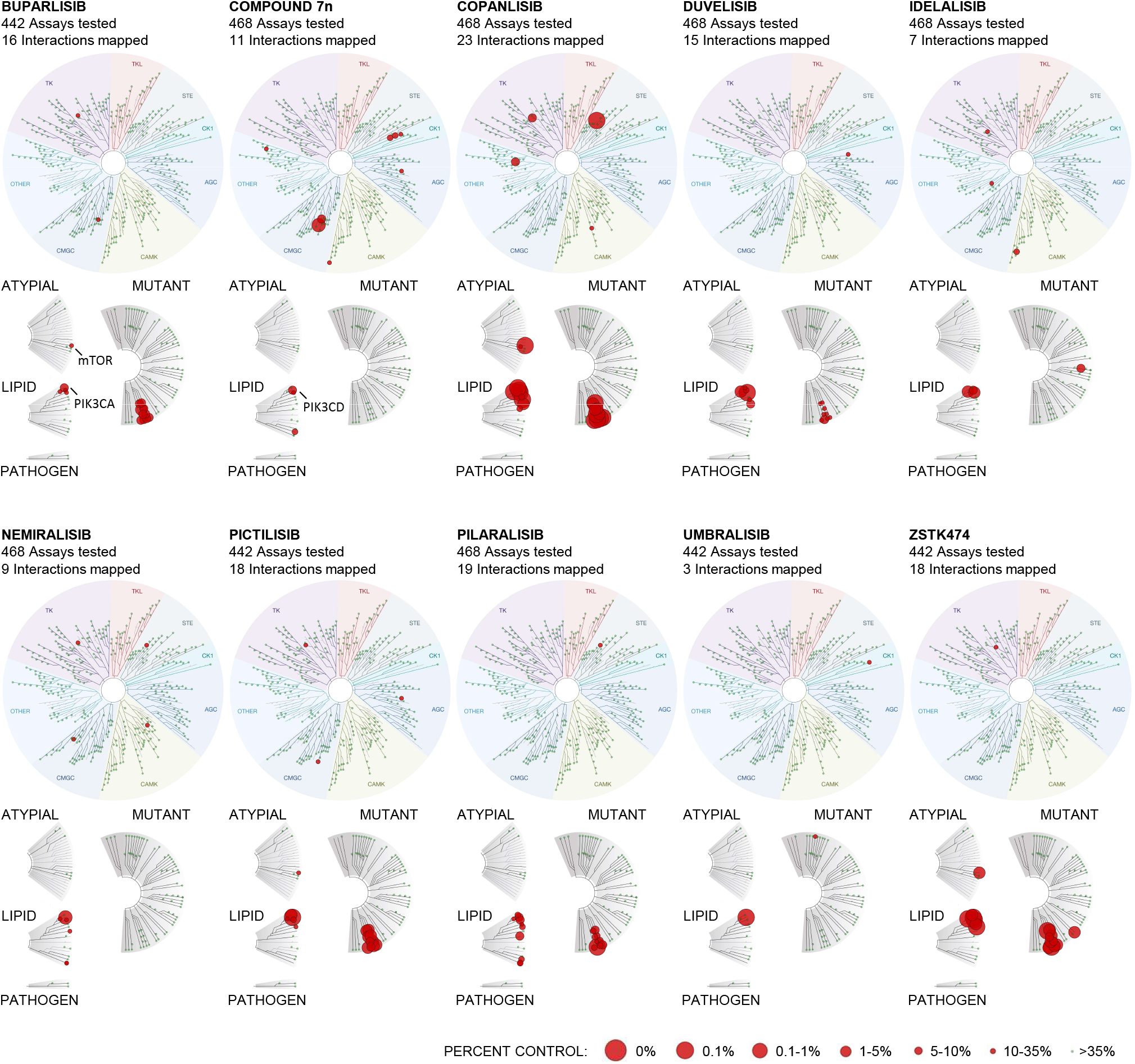
Target specificity and activity profile for ten PI3Ki. The activity of buparlisib, compound 7n, copanlisib, duvelisib, idelalisib, nemiralisib, pictilisib, pilaralisib, umbralisib, and ZSTK474 were profiled at 1 μM over a panel of up to 468 human kinases, including atypical, mutant, lipid, and pathogen kinases (lower dendrograms), using the DiscoveRx kinase assays. The upper dendrograms show (moving clockwise from the upper, purple section) tyrosine kinases (TK), tyrosine kinase-like kinases (TKL), STE protein kinases (STE), casein kinase 1 family (CK1), protein kinase A, G, and C families (AGC), Ca^2+^/calmodulin-dependent kinases (CAMK), CMGC kinase group (CMGC), and other. The size of the circles represents the percentage of target inhibition, with larger circles indicating a stronger inhibition compared with control, as defined in the scale.

**Figure 2.**
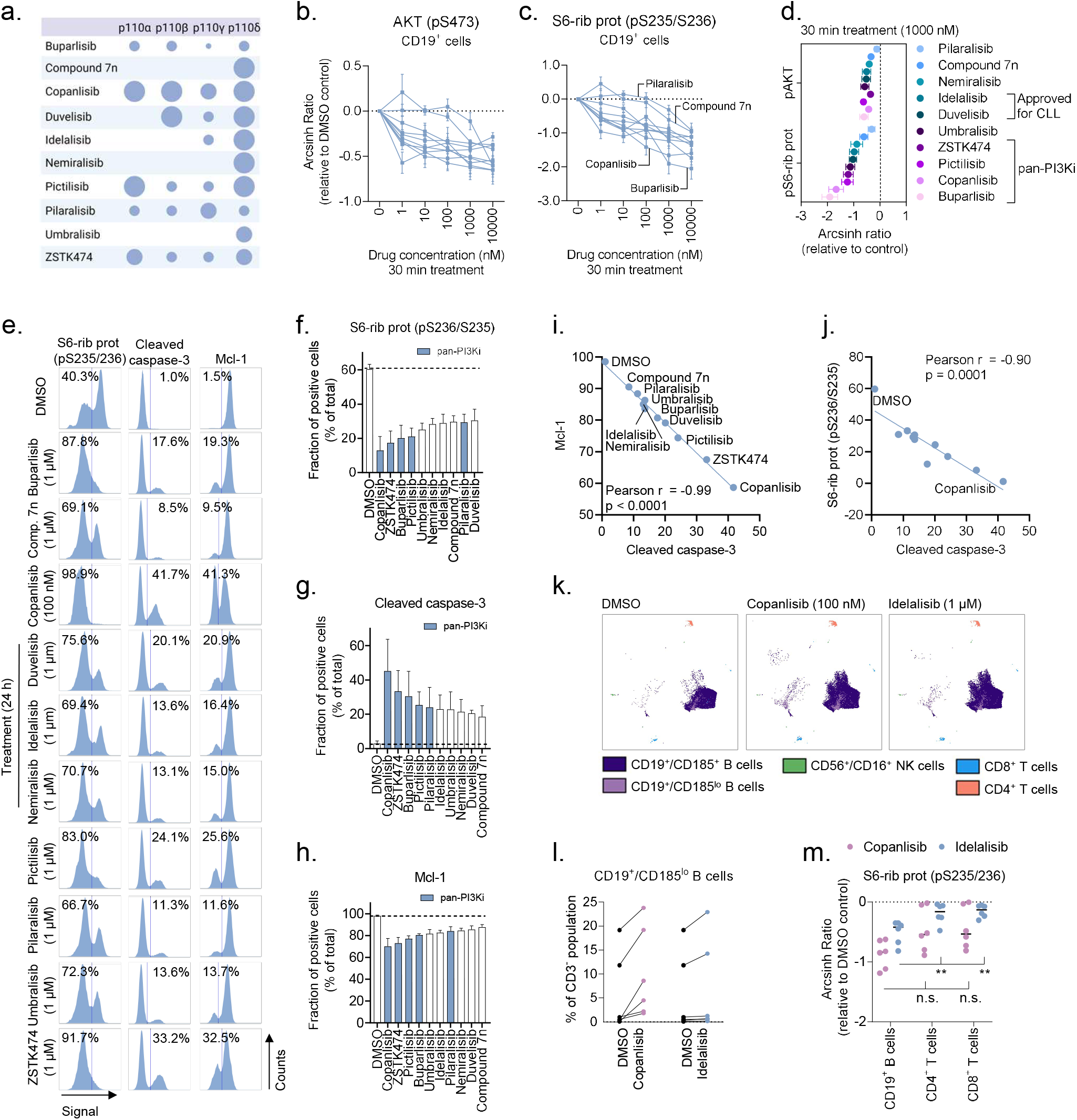
Drug-induced changes in cell signaling and cell viability are PI3Ki specific. a) Relative activity of the ten PI3Ki against p110 isoforms. The size of the circles represents the relative extent of inhibition, with larger circles indicating stronger inhibition. b-c) Peripheral blood mononuclear cells (PBMCs) from CLL patient samples (n=4) were treated with ten PI3Ki (Table 1) at the indicated concentrations for 30 min, followed by 5 min anti-IgM stimulation. The cells were then fixed, permeabilized and stained with the indicated antibodies. Signals were analyzed by flow cytometry. Results are shown for CD19^+^ B cells. Raw data were transformed to an arcsinh ratio relative to the signal in DMSO treated control cells, which was set to zero. Curves show the mean of the four experiments. Each curve represents the response to one inhibitor. Error bars indicate standard error of the mean (SEM). d) Results are shown for experiments described in (b-c). The mean inhibitory effects of the ten PI3Ki on phosphorylation of AKT (pS473) and S6-ribosomal protein (pS235/S236) are plotted at the 1000 nM concentrations. Error bars indicate SEM. e) CLL cells were simultaneously co-cultured with APRIL/BAFF/CD40L^+^ fibroblasts and treated with the indicated concentration of a PI3Ki or DMSO (0.1%) for 24h. The CLL cells were then separated from the fibroblast layer, fixed, permeabilized and stained with the indicated antibodies. Signals were analyzed by flow cytometry. The cells were gated on CD19. The numbers indicate the fraction (%) of cells in the respective gate. Results are shown for one representative experiment f-h) Experiments were performed as described in (e) on CLL cells from n=3 patients. The bars show the mean fraction (%) of positive cells ± SEM. The dotted line indicates the fraction of positive cells in the DMSO control. The blue bars indicate pan-PI3Ki. i-j) Pearson’s correlation analyses were performed on the indicated protein levels detected in (e). Each point represents one treatment. k) PBMCs from n=6 treatment naïve CLL patients were co-cultured with APRIL/BAFF/CD40L^+^ fibroblasts and DMSO (0.1%), copanlisib (100 nM), or idelalisib (1 µM) for 24h. The PBMCs were then separated from the fibroblast layer, fixed, barcoded, permeabilized and stained with surface markers and anti-S6-ribosomal protein (pS235/S236) antibody. Experiments were analyzed with a BD FACSymphony A5 cytometer (BD Biosciences) and further processed in Cytobank (https://cellmass.cytobank.org/cytobank/). FlowSOM clustering algorithm was applied to identify cell populations, which were validated by manual gating. The UMAP dimensionality reduction algorithm was used to visualize the data. The UMAP for one representative patient sample is shown. l) Experiments were performed as described in (k). CD19^+^/CD185^lo^ B cells were quantified as percent of CD3^-^ lymphocytes. m) Experiments were performed as described in (k). Signals are shown for CD19^+^ B cells, CD4^+^ T cells, and CD8^+^ T cells. Raw data were transformed to an arcsinh ratio relative to the signal in DMSO treated control cells, which was set to zero. Statistical testing was done with 2-way ANOVA. **p<0.01, n.s; not significant.

### Drug-induced changes in cell signaling are PI3Ki specific

To study how the different compounds inhibit PI3K signaling, CLL cells were treated with the individual PI3Ki at five concentrations (1 nM - 10000 nM) for 30 min. Copanlisib was tested at 10-fold lower concentrations (0.1 nM – 1000 nM) due to its higher potency.^38^ The cells were then stimulated with anti-IgM for 5 min to activate the BCR. As shown in **Figure 2b-c**, each PI3Ki inhibited the phosphorylation of downstream effectors AKT (pS473) and S6-ribosomal protein (pS235/S236), but with varying potency. Overall, the pan-PI3Ki buparlisib, copanlisib, pictilisib and ZSTK474 were most effective at inhibiting PI3K signaling (**Figure 2d**). To study whether this trend could be reproduced with longer incubation times, CLL cells were incubated with each PI3Ki for 24h. To prevent induction of spontaneous apoptosis, which would influence the read-out, the CLL cells were simultaneously co-cultured with fibroblasts expressing APRIL/BAFF/CD40L. As shown in **Figure 2e-f**, 24h incubation with a PI3Ki inhibited phosphorylation of S6-ribosomal protein (pS235/S236), and the pan-PI3Ki were again most effective. Furthermore, 24h incubation with a PI3Ki induced apoptosis, as indicated by increased expression of cleaved caspase-3 (**Figure 2e, g**). Similarly, expression of the anti-apoptotic protein Mcl-1 was reduced in response to PI3Ki exposure. These phenotypes were most pronounced in response to a pan-PI3Ki (**Figure 2g-h**). The drug-induced changes in cell signaling and protein expression showed a highly significant correlation (**Figure 2i-j**), indicating that inhibition of cell signaling can serve as a substitute marker for induction of apoptosis.

### Effects of copanlisib and idelalisib on different immune cell compartments

To study the effect of PI3K inhibition on different immune cell compartments, PBMCs from treatment naïve CLL patients were simultaneously cultured with fibroblasts expressing APRIL/BAFF/CD40L and treated either with DMSO (control), copanlisib (100 nM), or idelalisib (1 µM) for 24h (**Figure 2k**). The CD185^lo^ (CXCR5) B cell population was shown to consistently expand in response to copanlisib treatment (**Figure 2l**). This effect was not as pronounced upon idelalisib treatment, although two of the patient samples showed a similar trend (**Figure 2l**). CD185 is a mediator of CLL tissue homing, and it has been shown to decrease also in response to ibrutinib + rituximab treatment.^39^ Other analyzed immune subsets were not significantly modulated by the treatments. This is in agreement with a previous study showing specificity of idelalisib and copanlisib towards CLL B cells.^38^ However, we observed that copanlisib, but not idelalisib, reduced S6-ribosomal protein phosphorylation also in CD4^+^ and CD8^+^ T cells (**Figure 2m**).

### Pan-PI3Ki show activity in idelalisib-resistant lymphoma cells and idelalisib-refractory/intolerant CLL cells

To study the ability of the different PI3Ki to inhibit cell viability, 72h drug sensitivity assays were performed on the lymphoma cell lines VL51 and KARPAS1718 (**Figure 3a-b**). Each PI3Ki was tested at five different concentrations (1 nM - 10000 nM). Due to its higher potency, copanlisib was tested at 10-fold lower concentrations (0.1 nM - 1000 nM). In agreement with the effects observed on cell signaling, the pan-PI3Ki buparlisib, copanlisib, pictilisib and ZSTK474 were most effective at inhibiting cell viability in the parental cell lines (**Figure 3a-b**, black bars).

**Figure 3.**
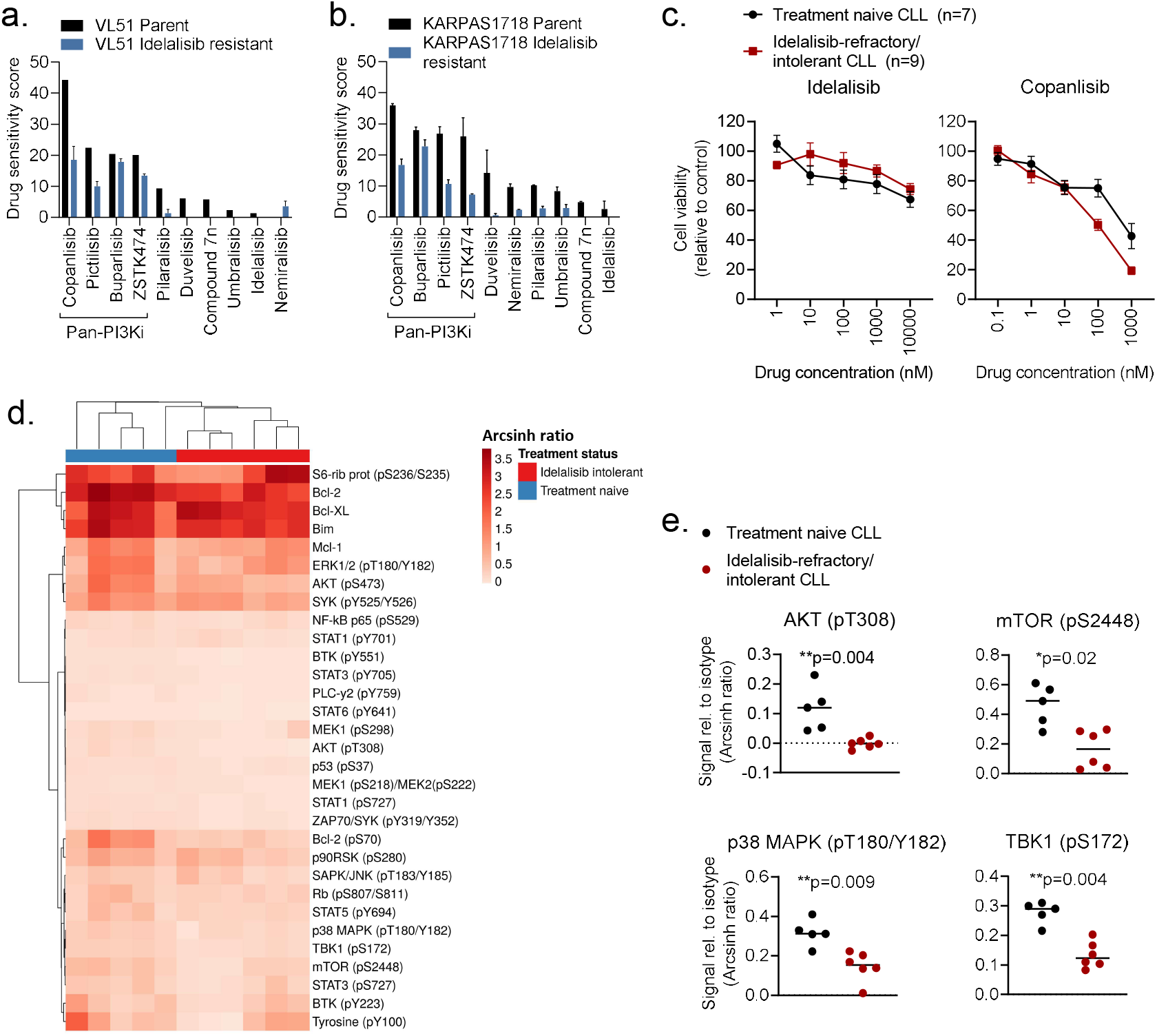
Pan-PI3Ki are active in idelalisib-refractory/intolerant CLL cells. a-b) VL51 and KARPAS1718 (parent and idelalisib resistant) cell lines were treated with the indicated PI3Ki at 5 concentrations (0.1 nM – 1000 nM for copanlisib, 1 nM – 10000 nM for the others) for 72h. Cell viability was assessed with the CellTiter-Glo assay. The drug sensitivity score was calculated for each treatment based on the area under the dose-response curve. High score indicates high sensitivity to the treatment. The experiment were performed twice or once (VL51 parent). The bars show average with standard deviation. c) PBMCs from treatment naïve CLL patients (n=7) or idelalisib refractory patients (n=9) were co-cultured with APRIL/BAFF/CD40L^+^ fibroblasts for 24h. The CLL cells were then separated from the fibroblast layer and treated with the indicated compound and concentrations for 72h. Cell viability was assessed with the CellTiter-Glo assay. The graphs show mean relative cell viability ± SEM. d) Freshly thawed PBMCs from treatment naïve CLL patients (n=5) or idelalisib refractory patients (n=6) were fixed, permeabilized and stained with antibodies against the indicated proteins (rows). Signals were detected in CD19^+^ B cells by flow cytometry. Raw data were transformed to an arcsinh ratio relative to the signal of an isotype control (color key), which was set to zero. The heatmap was created using ClustVis (https://biit.cs.us.ee/clustvis/). Rows are clustered using Manhattan distance and Ward linkage. Columns were clustered using correlation distance and average linkage. e) Phosphorylation levels of the indicated proteins are shown for experiments described in (d).The horizontal line indicates median. Statistical testing was done with the Mann-Whitney test.

To identify potential alternative therapies for patients who experience intolerance or develop resistance to idelalisib, we investigated whether idelalisib-resistant lymphoma cell lines remained sensitivity to other PI3Ki. As expected, PI3Ki showed reduced efficacy in idelalisib-resistant cells, but pan-PI3Ki still maintained a relatively high activity (**Figure 3a-b**). In the idelalisib resistant VL51 cell line, the activity of the four indicated pan-PI3Ki remained between 42% (copanlisib) and 87.5% (buparlisib) relative to the activity in the parent cell line. For KARPAS1718, the corresponding range was between 27.8% (ZSTK474) and 81.4% (buparlisib). The sensitivity to idelalisib was reduced in CLL cells from idelalisib-refractory/intolerant patients as well (**Figure 3c**, left). However, the sensitivity to copanlisib was maintained at similar or even higher levels than in CLL cells from treatment naïve patients (**Figure 3c**, right). This trend was also observed for the pan-PI3Ki buparlisib, pictilisib, and ZSTK474 (**Supplementary Figure 1**). These findings demonstrate that pan-PI3Ki remain active in idelalisib-refractory/intolerant cells.

### Idelalisib-refractory/intolerant CLL cells show reduced protein phosphorylation levels

To better understand the mechanisms underlying idelalisib-refractory/intolerant disease, we performed single-cell protein profiling of CLL cells from patients who were either treatment naïve or refractory/intolerant to idelalisib (**Figure 3d**). We found that expression or phosphorylation of several proteins was downregulated in CLL cells from idelalisib-refractory/intolerant patients relative to treatment naïve patients, including Bcl-2 (p=0.02 with an unpaired t test), Bcl-2 (pS70) (p=0-009), BTK (pY551) (p=0.004), MEK1 (pS218)/MEK2 (pS222) (p=0.004), p53 (pS37) (p=0.004), PLCγ2 (pY759) (p=0.02), STAT1 (pS727) (p=0.01), STAT3 (pY705) (p=0.02), STAT6 (pY641) (p=0.004), and ZAP70/SYK (pY319/Y352) (p=0.004). Of note, phosphorylation of proteins downstream of PI3K, including AKT (pT308), mTOR (pS2448) and TBK1 (pS172), were all significantly reduced in idelalisib-refractory/intolerant CLL cells (**Figure 3e**). These findings support previous reports that have demonstrated that higher response rates to copanlisib are associated with high expression of PI3K/BCR signaling pathway genes.^40;41^ We also found that the activity of p38 MAPK (pT180/Y182) was reduced in idelalisib-refractory/intolerant cells (**Figure 3e**). Interestingly, we previously showed that low phosphorylation levels of p38 MAPK correlates with poor response to venetoclax,^18^ suggesting that cell signaling profiles may provide response markers for different classes of targeted therapies.

### PI3Ki act in synergy with venetoclax to induce apoptosis in CLL cells

Targeted therapies are increasingly studied in combinations, and venetoclax has been reported to be a good combination partner for copanlisib in B cell lymphoma (NCT03886649).^42^ We therefore investigated the potential benefit of combining a PI3Ki with the Bcl-2 antagonist venetoclax. CLL cells were treated with five different concentrations of each PI3Ki, venetoclax, or their combinations, for 30 min. The expression level of cleaved caspase-3 was then analyzed by flow cytometry. None of the PI3Ki as a single agent induced apoptosis after this short incubation time, while treatment with venetoclax did (**Figure 4a**). Interestingly, each PI3Ki + venetoclax combination induced higher levels of cleaved caspase-3 than the venetoclax treatment alone (**Figure 4a**), suggesting a synergistic effect of the combination. Dose-response combination assays confirmed synergy among all ten combinations (**Figure 4b-c**).

**Figure 4.**
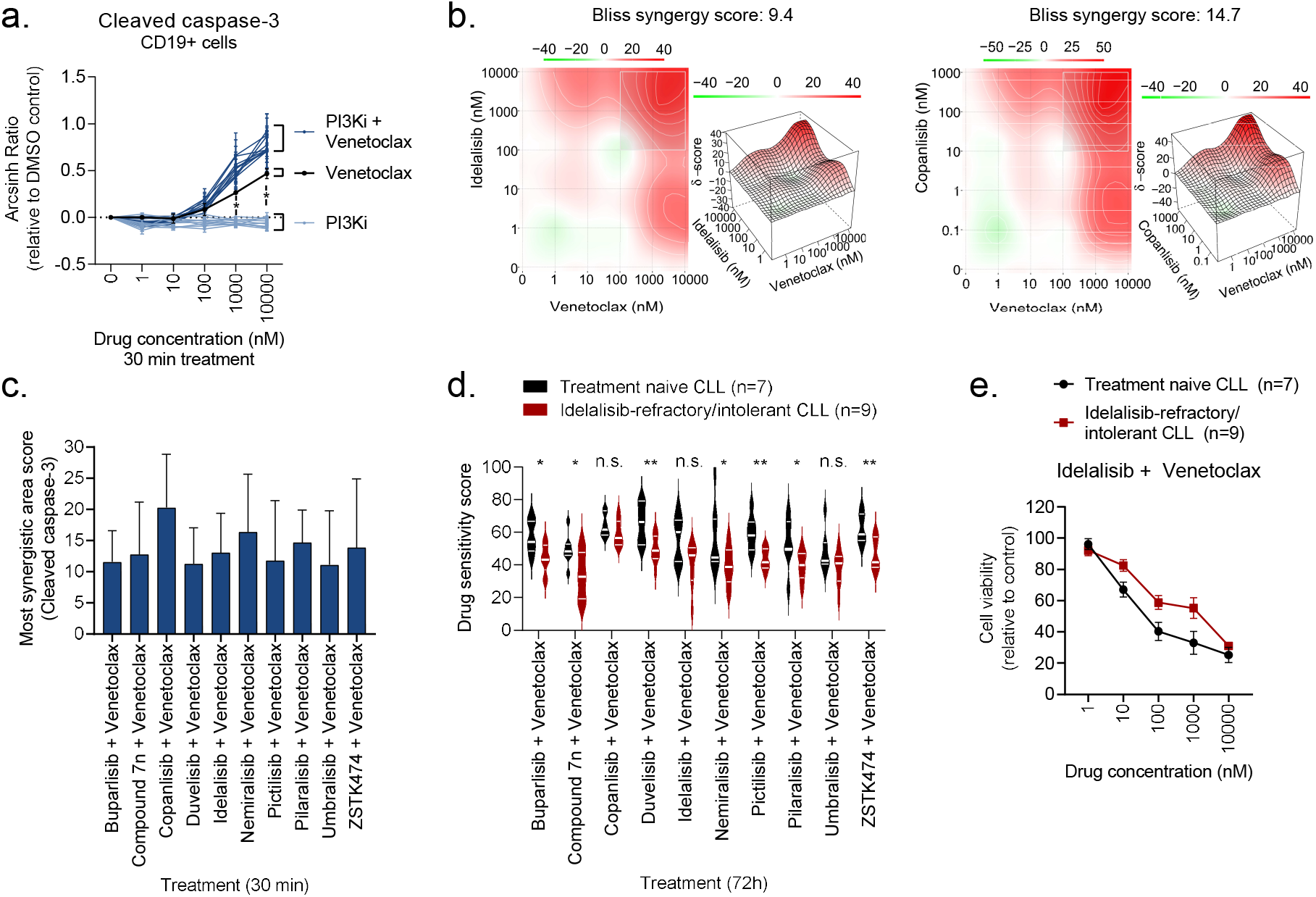
PI3Ki act in synergy with venetoclax. a) PBMCs from CLL patient samples (n=4) were treated with a PI3Ki, venetoclax, or PI3Ki + venetoclax combinations, at the indicated concentrations for 30 min, followed by 5 min anti-IgM stimulation. The cells were then fixed, permeabilized and stained with anti-cleaved caspase-3. Signals were analyzed by flow cytometry. Results are shown for CD19^+^ B cells. Raw data were transformed to an arcsinh ratio relative to the signal in DMSO treated control cells, which was set to zero. Curves show the mean ± SEM. Statistical testing was done with a one-way ANOVA with Holm-Sidak’s multiple comparisons test. *p<0.05. b) Normalized data from experiments described in (a) were used in DECREASE (https://decrease.fimm.fi/) to predict the full drug combination dose-response matrices, which were analyzed using SynergyFinder (https://synergyfinder.fimm.fi/) to score the synergy of the drug combinations. Bliss synergy over the full matrix is indicated. A representative plot is shown for the idelalisib + venetoclax and copanlisib + venetoclax combinations. c) Results are shown for analyses described in (b). The most synergistic area score was calculated by SynergyFinder (https://synergyfinder.fimm.fi/) for the indicated combination treatments. Bars show mean (n=4) ± SEM. d) PBMCs from treatment naïve CLL patients (n=7) or idelalisib-refractory/intolerant patients (n=9) were co-cultured with APRIL/BAFF/CD40L^+^ fibroblasts for 24h. The CLL cells were then separated from the fibroblast layer and treated with the indicated drug combinations for 72h. Cell viability was assessed with the CellTiter-Glo assay. The drug sensitivity score was calculated for each treatment based on the area under the dose-response curve. High score indicates high sensitivity to the treatment. Violin plots show min to max response with lines at quartiles and median. Statistical testing was done with an unpaired t test. *p<0.05, **p<0.01, n.s; not significant. e) PBMCs from treatment naïve CLL patients (n=7) or idelalisib-refractory/intolerant patients (n=9) were co-cultured with APRIL/BAFF/CD40L^+^ fibroblasts for 24h. The CLL cells were then separated from the fibroblast layer and treated with idelalisib + venetoclax combination at the indicated concentrations for 72h. Cell viability was assessed with the CellTiter-Glo assay. The graph shows mean relative cell viability ± SEM.

Next, we tested whether PI3Ki + venetoclax combinations were effective in co-inhibiting the viability of CLL cells from idelalisib-refractory/intolerant patients over 72h. Overall, the sensitivities to most of these combinations were reduced in idelalisib-refractory/intolerant CLL relative to treatment naïve CLL (**Figure 4d**). However, the combinations did reduce the cell viability in a concentration-dependent manner consistently in both patient groups, as shown for idelalisib + venetoclax (**Figure 4e**). Taken together, these findings show that PI3Ki act in synergy with venetoclax to induce apoptosis, and that PI3Ki + venetoclax combinations are active in both treatment naïve and idelalisib-refractory/intolerant CLL.

### *Ex vivo* drug sensitivity predicts clinical response in a CLL index patient

A 64-year-old, female CLL patient (CLL150, **Supplementary Table 4**) with unmutated IGVH (UM-CLL) and TP53 mutation presented with refractory disease after sequential treatment with FCR, FC, ibrutinib, idelalisib, and venetoclax (**Figure 5a**). To guide the next treatment for this patient, we collected a blood sample when her disease was progressing on venetoclax (T1, **Figure 5a**), and performed a drug sensitivity screen on her PBMCs. We found that the patient cells were sensitive to combined PI3Ki + venetoclax treatment (**Figure 5b**). The combination of idelalisib + venetoclax was selected as next treatment since these therapies are available in Norway (**Figure 5b**, red curve). The patient obtained a partial response, and the blood values nearly normalized while on treatment (**Figure 5c**). Since the patient suffered intolerable gastrointestinal adverse effects, the treatment was stopped after three months. Nevertheless, this case suggests that *ex vivo* drug sensitivity can predict clinical treatment response, and that CLL which is refractory to idelalisib and venetoclax monotherapies may still respond to idelalisib + venetoclax combination therapy.

**Figure 5.**
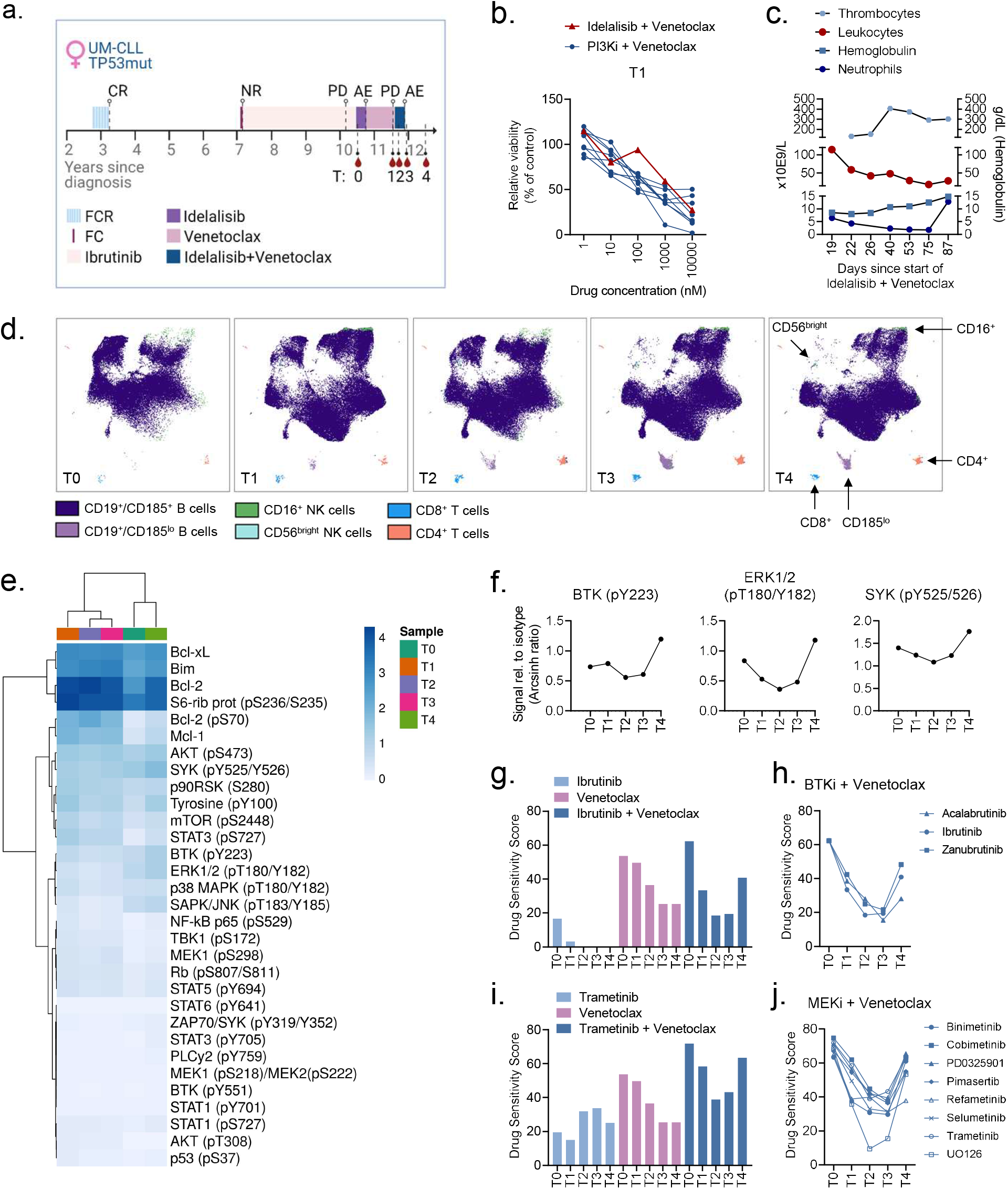
*Ex vivo* drug sensitivity predicts clinical response in a CLL index patient. a) Time-line illustrating the treatment history and disease progression of patient CLL150. T0-T4 indicate the time-points when blood samples were collected from the patient. AE, adverse event; CR, complete response; FC(R), fludarabine + cyclophosphamide (+ rituximab); NR, no response; PD, progressive disease. b) PBMCs from sample T1 were co-cultured with APRIL/BAFF/CD40L^+^ fibroblasts for 24h. The CLL cells were then separated from the fibroblast layer and treated with PI3Ki + venetoclax combination at the indicated concentrations for 72h. Cell viability was assessed with the CellTiter-Glo assay. c) Blood counts of CLL150 in response to treatment with idelalisib + venetoclax combination. Note the non-linear time scale. d) PBMCs from CLL150 T0-T4 were fixed, barcoded, permeabilized and stained with surface markers. Experiments were analyzed with a BD FACSymphony A5 cytometer (BD Biosciences) and further processed in Cytobank (https://cellmass.cytobank.org/cytobank/). At each time-point, 70.000 events were collected. FlowSOM clustering algorithm was applied to identify cell populations, which were validated by manual gating. The UMAP dimensionality reduction algorithm was used to visualize the data. Immune cell compartments are indicated in the color key. e) Freshly thawed PBMCs from T0-T4 were fixed, permeabilized and stained with antibodies against the indicated proteins (rows). Signals were detected in CD19^+^ B cells by flow cytometry. Raw data were transformed to an arcsinh ratio relative to the signal of an isotype control, which was set to zero. The heatmap was created using ClustVis (https://biit.cs.us.ee/clustvis/). Both rows and columns were clustered using Manhattan distance and Ward linkage algorithm. f) Phosphorylation levels of the indicated proteins are shown for experiments described in (d). g-j) PBMCs from T0-T4 were co-cultured with APRIL/BAFF/CD40L^+^ fibroblasts for 24h. The CLL cells were then separated from the fibroblast layer and treated with the indicated drugs for 72h. Cell viability was assessed with the CellTiter-Glo assay. The drug sensitivity score (DSS) was calculated for each treatment based on the area under the dose-response curve. High score indicates high sensitivity to the treatment.

To study the evolution of the patient’s disease, we mapped the immune cell phenotype, cell signaling, and protein expression landscapes in PBMCs collected at five different sampling times: while the patient was on treatment with idelalisib (T0), venetoclax (T1), or idelalisib + venetoclax (T2), and after the end of treatment (T3, T4) (**Figure 5a**). The B cell population was dominating at all time-points, with a small increase (5%) in the CD185^lo^ (CXCR5) population at T3 (**Figure 5d**). Relative numbers of CD4^+^ T cells increased from 28.6% of the CD3^+^ population at T2 to 55% at T3, but were reduced to 48.9% at T4 (**Figure 5d**). CD56^+^CD16^+^ NK cells showed a similar trend, increasing from 13% of the CD3^-^ CD19^-^ population at T2 to 30% at T3, then down to 11% at T4. Reshaping of immune cell compartments in response to venetoclax treatment has been reported by us and others.^25;43^ The modest reshaping of the immune phenotype at T3 was in agreement with the partial clinical response to idelalisib + venetoclax therapy.

We further observed that B-cell signaling was reduced by the combination therapy (T2), but it increased again when the therapy was stopped (T3, T4) (**Figure 5e-f**). Increased phosphorylation of Bruton’s tyrosine kinase (BTK) and extracellular signal-regulated kinase (ERK) at T4 was mirrored by increased sensitivity to venetoclax combined either with a BTK or MEK inhibitor (**Figure 5g-j**). Unfortunately, the patient passed away and the *in vivo* efficacy of these alternative combination therapies could not be tested.

### *Ex vivo* drug sensitivity stratifies long-term responders to idelalisib therapy

To more systematically assess the predictive value of *ex vivo* drug sensitivity profiles, we analyzed drug responses to 73 combinations (**Supplementary Table 2**) on CLL cells from patients treated with idelalisib (“JB” samples, **Supplementary Table 4**). CLL cells collected at baseline from patients who obtained a long-term response to idelalisib showed significantly higher drug sensitivity scores than CLL cells from patients who obtained a short-term response (i.e., patients who developed resistance to idelalisib) (**Figure 6a**). Annotated drug responses are shown in **Supplementary Figure 2**. After correcting for multiple comparisons, we identified significant differences in responses to three PI3Ki + venetoclax combinations (**Figure 6b**). Interestingly, cells collected at the time the patients were responding to idelalisib showed lower levels of AKT (pS473) in long-term responders than in short-term responders (**Figure 6c**). However, this difference was not statistically significant, possible due to the low number of samples in each group (n=3). Taken together, these results demonstrate that functional profiling data can identify long-term responders to idelalisib, and warrant further studies in larger cohorts and clinical trials.

**Figure 6.**
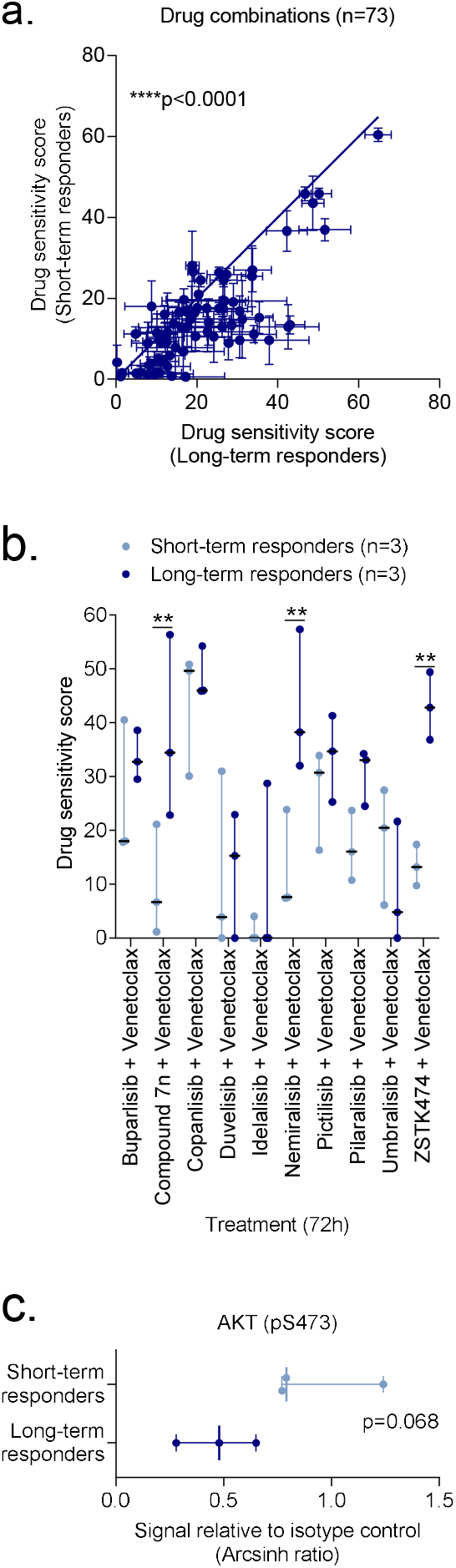
*Ex vivo* drug sensitivity stratifies responders to idelalisib. a) Drug sensitivity screens were performed with 73 drug combinations on PBMCs collected from CLL patients before the patients started treatment with idelalisib (n=3 short-term responders, i.e. developed resistance to idelalisib, and n=3 long-term responders). Each data point indicates the mean drug sensitivity score to one drug combination. Error bars show standard error of the mean (SEM) over the patients (n=3). The diagonal line indicates equal sensitivity in long- and short-term responders. Statistical testing was done with a paired t test. ****p<0.0001. b) As in (a), but with the indicated PI3Ki + venetoclax combinations. The dot plot shows median with range. Statistical testing was done with a 2-way ANOVA with Sidak’s multiple comparisons within treatment groups for the 73 drug combinations. **p<0.01. c) PBMCs collected from short-term and long-term responders at the time of response to idelalisib were fixed, permeabilized and stained with anti-AKT (pS473). Signals were detected in CD19^+^ B cells by flow cytometry. Raw data were transformed to an arcsinh ratio relative to the signal of an isotype control, which was set to zero. The scatter dot plot shows median with range. Statistical testing was done with an unpaired t test.

## DISCUSSION

Targeted therapies have considerably improved patient outcome in CLL, but development of treatment intolerance and resistance remains to be clinical challenges. One strategy to prevent resistance to monotherapy is to combine therapies.^8^ Here, we studied the efficacy, specificity, and synergy of ten PI3Ki, both as single agents and in combination with the Bcl-2 antagonist venetoclax. Venetoclax targets the intrinsic apoptotic pathway and is therefore an attractive partner for BCR inhibitors. The combination of venetoclax with the BTK inhibitor ibrutinib has shown promising results in CLL.^44;45^ Numerous additional studies on CLL are currently investigating the effect of venetoclax in combination with other targeted therapies, including various PI3Ki (NCT03534323, NCT03379051, NCT03801525). Resistance to both PI3Ki and venetoclax may be prevented or delayed with this strategy, as a suggested mechanism of resistance to venetoclax is upregulation of the PI3K/AKT/mTOR pathway.^46^

Here, we showed that pan-PI3Ki were more effective than p110δ selective inhibitors at inducing death of CLL cells, both as single agents and in combination with venetoclax.

Studies of copanlisib in relapsed or refractory lymphoma have demonstrated significant efficacy and a manageable safety profile,^40;47;48^ suggesting that copanlisib is a relevant treatment option for lymphoproliferative diseases. Copanlisib plus venetoclax combination showed the highest efficacy in our *ex vivo* assays. Synergy between these compounds has been demonstrated in B- and T-cell lymphoma models,^42^ and the combination is currently being studied in relapsed/refractory B-cell lymphomas (NCT03886649). The results will provide further information on its safety and efficacy.

Other strategies to prevent treatment resistance include improved patient stratification or precision medicine.^8^ CLL is a highly heterogeneous disease, and better model systems and biomarkers to guide clinical decision-making are likely to be useful. Functional assays can be valuable to this end.^2;26^ *Ex vivo* drug sensitivity has successfully predicted clinical activity in hematological malignancies.^49-51^ For instance, the EXALT trial (NCT03096821) investigated the feasibility and clinical impact of image-based *ex vivo* drug sensitivity-guided treatment decisions in patients with aggressive refractory hematological malignancies.^50^ The study showed that integration of sensitivity testing in clinical decisions led to improved treatment outcomes.^50^

Here, we showed that *ex vivo* drug screening accurately predicted drug response in a monotherapy refractory CLL patient. *Ex vivo* drug sensitivities could further stratify patients based on clinical treatment responses, suggesting that *ex vivo* drug screens can serve as a predictive biomarker for treatment outcome. Our analyses showed that CLL cells from patients who are resistant to idelalisib remain sensitive to pan-PI3Ki. This finding is in agreement with studies showing that CLL patients who fail on a targeted therapy may still benefit from a second therapy in the same drug class,^20-22^ and warrants further studies on how to maximize the clinical value of PI3Ki.

Taken together, our findings indicate PI3Ki drug class activity in idelalisib-refractory/intolerant CLL and suggest that *ex vivo* drug sensitivity may guide precision medicine and predict treatment responses.

## ACKNOWLEDGEMENTS

The authors are thankful to all patients who contributed to this study. We are grateful to Silje Hjellbrekke, Hallvard Zapffe, Mentowa Fürst Bright and Martine Schrøder for technical assistance. We thank the High-Throughput Chemical Biology Screening Platform at Centre for Molecular Medicine Norway (NCMM), University of Oslo, and the High Throughput Biomedicine Unit at Institute for Molecular Medicine Finland (FIMM), University of Finland, for assistance with drug sensitivity screens. This work was supported by the Research Council of Norway under the frames of ERA PerMed (project number 322898) and Digital Life Norway (project number 294916), the Norwegian Cancer Society, the Regional Health Authority for South-Eastern Norway, Stiftelsen Kristian Gerhard Jebsen (Grant 19), Lilly Constance og Karl Ingolf Larssons stiftelse, and the Medical Student Research Program at the University of Oslo. F.B. was supported by Swiss National Science Foundation (SNSF 31003A_163232/1). J.R.B. was supported by NIH R01 CA 213442. T.A. was supported by the Norwegian Cancer Society, Radium Hospital Foundation and the Academy of Finland (grants 313267, 326238, and 344698).

## AUTHORSHIP

### Contribution

S.S.S. designed the study. Y.Y., L.K., A.U. and S.S.S. performed the experiments. Y.Y., L.K., P.A., A.U., T.A. and S.S.S. analyzed and interpreted the data with input from K.T., whereas I.M., S.M.F., A.K.H., A.R.M., E.N., J.R.B. and G.E.T. contributed with patient samples. A.J.A. and F.B. contributed with cell lines. S.S.S. wrote the article. All authors read and commented on the manuscript and approved the final version.

## SUPPLEMENT

**Supplementary Table 1**. Compound library

**Supplementary Table 2**. Drug combinations

**Supplementary Table 3**. Kinome scan data

**Supplementary Table 4**. Patient characteristics

B, bendamustine; C, cyclophosphamide; Chl; chlorambucil; f, female; F, fludarabine; m, male; M, IGVH mutated CLL; n.e., not established; OFA, ofatumumab; P, prednisone; R, rituximab; UM, IGVH unmutated CLL; V, vincristine

**Supplementary Figure 1. *Ex vivo* drug sensitivity to PI3Ki and venetoclax single agents and combinations in CLL cells**

PBMCs from the indicated CLL samples (n=27; columns) were co-cultured with APRIL/BAFF/CD40L^+^ fibroblasts for 24h. The CLL cells were then separated from the fibroblast layer and treated with the indicated compounds and combinations (n=21; rows) for 72h. Cell viability was assessed with the CellTiter-Glo assay. The drug sensitivity score (DSS; color key) was calculated for each treatment based on the modified area under the dose-response curve. High score indicates high sensitivity to the treatment. The heatmap was created using ClustVis (https://biit.cs.us.ee/clustvis/).

**Supplementary Figure 2. *Ex vivo* drug sensitivity stratifies responders to idelalisib**

Drug sensitivity screens were performed with the indicated 73 drug combinations on PBMCs collected from CLL patients before the patients started treatment with idelalisib (n=3 short-term responders, i.e. developed resistance to idelalisib, and n=3 long-term responders). Mean ± standard error of the mean (SEM) are shown.

