## Supplementary Table 2 for "Functional testing of PI3K inhibitors stratifies responders to idelalisib and identifies treatment vulnerabilities in idelalisib-refractory/intolerant chronic lymphocytic leukemia"

**Supplementary Table 2.** Drug combinations

| Drug 1 | Drug 2 |
|---|---|
| Acalabrutinib | AZD6738 |
| Acalabrutinib | Quizartinib |
| Alisertib | Crizotinib |
| Buparlisib | Acalabrutinib |
| Buparlisib | Ibrutinib |
| Buparlisib | Venetoclax |
| Compound 7n | Acalabrutinib |
| Compound 7n | Ibrutinib |
| Compound 7n | Venetoclax |
| Copanlisib | Acalabrutinib |
| Copanlisib | Ibrutinib |
| Copanlisib | Venetoclax |
| Cytarabine | Nutlin 3a |
| Duvelisib | Acalabrutinib |
| Duvelisib | Ibrutinib |
| Duvelisib | Venetoclax |
| Ibrutinib | Chlorambucil |
| Ibrutinib | Fludarabine |
| Ibrutinib | Quizartinib |
| Ibrutinib | Selinexor |
| Ibrutinib | SNX-5422 |
| Idelalisib | Acalabrutinib |
| Idelalisib | Ibrutinib |
| Idelalisib | JQ1 |
| Idelalisib | Quizartinib |
| Idelalisib | Ruxolitinib |
| Idelalisib | Trametinib |
| Idelalisib | Venetoclax |
| JQ1 | Palbociclib |
| JQ1 | Ruxolitinib |
| JQ1 | Sorafenib |
| Lenalidomide | Chlorambucil |
| Lenalidomide | Dexamethasone |
| Methylprednisolone | Lenalidomide |
| Nemiralisib | Acalabrutinib |
| Nemiralisib | Ibrutinib |
| Nemiralisib | Venetoclax |
| Palbociclib | Doramapimod |
| Palbociclib | Quizartinib |
| Palbociclib | Ruxolitinib |
| Palbociclib | Sorafenib |
| Panobinostat | Ruxolitinib |
| Pictilisib | Acalabrutinib |
| Pictilisib | Ibrutinib |

| | |
|---|---|
| Pictilisib | Venetoclax |
| Pilaralisib | Acalabrutinib |
| Pilaralisib | Ibrutinib |
| Pilaralisib | Venetoclax |
| Ruxolitinib | Cabozantinib |
| Trametinib | Palbociclib |
| Umbralisib | Acalabrutinib |
| Umbralisib | Ibrutinib |
| Umbralisib | Venetoclax |
| Valproic acid | 2-chlorodeoxyadenosine |
| Vandetanib | Vemurafenib |
| Venetoclax | Acalabrutinib |
| Venetoclax | Binimetinib |
| Venetoclax | Cobimetinib |
| Venetoclax | Dasatinib |
| Venetoclax | Ibrutinib |
| Venetoclax | Ibrutinib |
| Venetoclax | Palbociclib |
| Venetoclax | PD0325901 |
| Venetoclax | Pimasertib |
| Venetoclax | Refametinib |
| Venetoclax | Ruxolitinib |
| Venetoclax | Selumetinib |
| Venetoclax | Sorafenib |
| Venetoclax | Trametinib |
| Venetoclax | U0126 |
| ZSTK474 | Acalabrutinib |
| ZSTK474 | Ibrutinib |
| Acalabrutinib | AZD6738 |
| Acalabrutinib | Quizartinib |
| Alisertib | Crizotinib |
| Buparlisib | Acalabrutinib |
| Buparlisib | Ibrutinib |
| Buparlisib | Venetoclax |
| Compound 7n | Acalabrutinib |
| Compound 7n | Ibrutinib |
| Compound 7n | Venetoclax |
| Copanlisib | Acalabrutinib |
| Copanlisib | Ibrutinib |
| Copanlisib | Venetoclax |
| Cytarabine | Nutlin 3a |
| Duvelisib | Acalabrutinib |
| Duvelisib | Ibrutinib |
| Duvelisib | Venetoclax |
| Ibrutinib | Chlorambucil |
| Ibrutinib | Fludarabine |

| | |
|---|---|
| Ibrutinib | Quizartinib |
| Ibrutinib | Selinexor |
| Ibrutinib | SNX-5422 |
| Idelalisib | Acalabrutinib |
| Idelalisib | Ibrutinib |
| Idelalisib | JQ1 |
| Idelalisib | Quizartinib |
| Idelalisib | Ruxolitinib |
| Idelalisib | Trametinib |
| Idelalisib | Venetoclax |
| JQ1 | Palbociclib |
| JQ1 | Ruxolitinib |
| JQ1 | Sorafenib |
| Lenalidomide | Chlorambucil |
| Lenalidomide | Dexamethasone |
| Methylprednisolone | Lenalidomide |
| Nemiralisib | Acalabrutinib |
| Nemiralisib | Ibrutinib |
| Nemiralisib | Venetoclax |
| Palbociclib | Doramapimod |
| Palbociclib | Quizartinib |
| Palbociclib | Ruxolitinib |
| Palbociclib | Sorafenib |
| Panobinostat | Ruxolitinib |
| Pictilisib | Acalabrutinib |
| Pictilisib | Ibrutinib |
| Pictilisib | Venetoclax |
| Pilaralisib | Acalabrutinib |
| Pilaralisib | Ibrutinib |
| Pilaralisib | Venetoclax |
| Ruxolitinib | Cabozantinib |
| Trametinib | Palbociclib |
| Umbralisib | Acalabrutinib |
| Umbralisib | Ibrutinib |
| Umbralisib | Venetoclax |
| Valproic acid | 2-chlorodeoxyadenosine |
| Vandetanib | Vemurafenib |
| Venetoclax | Acalabrutinib |
| Venetoclax | Binimetinib |
| Venetoclax | Cobimetinib |
| Venetoclax | Dasatinib |
| Venetoclax | Ibrutinib |
| Venetoclax | Ibrutinib |
| Venetoclax | Palbociclib |
| Venetoclax | PD0325901 |
| Venetoclax | Pimasertib |

| | |
|---|---|
| Venetoclax | Refametinib |
| Venetoclax | Ruxolitinib |
| Venetoclax | Selumetinib |
| Venetoclax | Sorafenib |
| Venetoclax | Trametinib |
| Venetoclax | U0126 |
| ZSTK474 | Acalabrutinib |
| ZSTK474 | Ibrutinib |
| ZSTK474 | Venetoclax |
