## Supplementary Table 3 for "Functional testing of PI3K inhibitors stratifies responders to idelalisib and identifies treatment vulnerabilities in idelalisib-refractory/intolerant chronic lymphocytic leukemia"

| Kinase target | Compound 7n | Copanlisib | Nemiralisib | Pilaralisib |
|---|---|---|---|---|
| AAK1 | 49 | 100 | 63 | 65 |
| ABL1(E255K)-phosphorylated | 83 | 72 | 52 | 80 |
| ABL1(F317I)-nonphosphorylated | 100 | 78 | 100 | 100 |
| ABL1(F317I)-phosphorylated | 92 | 100 | 99 | 99 |
| ABL1(F317L)-nonphosphorylated | 63 | 97 | 68 | 81 |
| ABL1(F317L)-phosphorylated | 97 | 100 | 81 | 91 |
| ABL1(H396P)-nonphosphorylated | 72 | 71 | 47 | 85 |
| ABL1(H396P)-phosphorylated | 96 | 76 | 81 | 100 |
| ABL1(M351T)-phosphorylated | 100 | 100 | 81 | 94 |
| ABL1(Q252H)-nonphosphorylated | 81 | 51 | 63 | 76 |
| ABL1(Q252H)-phosphorylated | 94 | 71 | 73 | 100 |
| ABL1(T315I)-nonphosphorylated | 87 | 100 | 89 | 98 |
| ABL1(T315I)-phosphorylated | 94 | 84 | 95 | 91 |
| ABL1(Y253F)-phosphorylated | 83 | 68 | 63 | 85 |
| ABL1-nonphosphorylated | 71 | 63 | 61 | 74 |
| ABL1-phosphorylated | 97 | 76 | 89 | 100 |
| ABL2 | 97 | 83 | 92 | 98 |
| ACVR1 | 97 | 71 | 72 | 100 |
| ACVR1B | 100 | 96 | 100 | 100 |
| ACVR2A | 100 | 95 | 100 | 100 |
| ACVR2B | 97 | 97 | 91 | 99 |
| ACVRL1 | 100 | 82 | 48 | 90 |
| ADCK3 | 73 | 76 | 100 | 100 |
| ADCK4 | 86 | 75 | 72 | 58 |
| AKT1 | 89 | 90 | 100 | 65 |
| AKT2 | 98 | 98 | 87 | 98 |
| AKT3 | 100 | 100 | 100 | 94 |
| ALK | 100 | 77 | 100 | 100 |
| ALK(C1156Y) | 96 | 80 | 89 | 89 |
| ALK(L1196M) | 95 | 82 | 98 | 94 |
| AMPK-alpha1 | 93 | 81 | 91 | 87 |
| AMPK-alpha2 | 98 | 95 | 97 | 79 |
| ANKK1 | 71 | 100 | 75 | 77 |
| ARK5 | 89 | 89 | 93 | 100 |
| ASK1 | 90 | 57 | 97 | 94 |
| ASK2 | 100 | 67 | 100 | 100 |
| AURKA | 92 | 90 | 94 | 98 |
| AURKB | 100 | 94 | 100 | 100 |
| AURKC | 34 | 98 | 81 | 100 |
| AXL | 88 | 84 | 80 | 97 |
| BIKE | 79 | 96 | 82 | 93 |
| BLK | 93 | 80 | 53 | 93 |
| BMPR1A | 100 | 88 | 100 | 100 |
| BMPR1B | 61 | 55 | 68 | 100 |
| BMPR2 | 88 | 86 | 91 | 100 |
| BMX | 97 | 85 | 83 | 98 |
| BRAF | 96 | 78 | 88 | 100 |
| BRAF(V600E) | 100 | 91 | 92 | 100 |
| BRK | 81 | 100 | 86 | 79 |
| BRSK1 | 94 | 94 | 48 | 100 |
| BRSK2 | 73 | 100 | 63 | 92 |
| BTK | 100 | 65 | 96 | 94 |
| BUB1 | 79 | 94 | 84 | 100 |
| CAMK1 | 45 | 89 | 82 | 90 |
| CAMK1B | 78 | 23 | 86 | 92 |
| CAMK1D | 72 | 89 | 86 | 95 |
| CAMK1G | 89 | 100 | 98 | 98 |
| CAMK2A | 90 | 100 | 87 | 94 |
| CAMK2B | 89 | 100 | 81 | 92 |
| CAMK2D | 93 | 80 | 100 | 96 |
| CAMK2G | 81 | 100 | 87 | 93 |
| CAMK4 | 100 | 81 | 100 | 100 |
| CAMKK1 | 95 | 100 | 99 | 98 |

| ≤ 20 | ≤ 2 |
|---|---|

| | | | | |
|---|---|---|---|---|
| CAMKK2 | 79 | 100 | 85 | 92 |
| CASK | 77 | 77 | 79 | 79 |
| CDC2L1 | 91 | 100 | 93 | 93 |
| CDC2L2 | 99 | 89 | 93 | 99 |
| CDC2L5 | 97 | 71 | 92 | 93 |
| CDK11 | 80 | 82 | 65 | 91 |
| CDK2 | 98 | 98 | 99 | 100 |
| CDK3 | 99 | 89 | 98 | 83 |
| CDK4 | 85 | 58 | 62 | 92 |
| CDK4-cyclinD1 | 92 | 72 | 13 | 100 |
| CDK4-cyclinD3 | 88 | 87 | 39 | 94 |
| CDK5 | 85 | 82 | 100 | 96 |
| CDK7 | 93 | 70 | 96 | 100 |
| CDK8 | 100 | 41 | 90 | 96 |
| CDK9 | 100 | 100 | 95 | 91 |
| CDKL1 | 71 | 90 | 88 | 81 |
| CDKL2 | 74 | 60 | 100 | 90 |
| CDKL3 | 93 | 81 | 81 | 99 |
| CDKL5 | 90 | 100 | 100 | 85 |
| CHEK1 | 97 | 99 | 93 | 88 |
| CHEK2 | 87 | 100 | 96 | 97 |
| CIT | 100 | 64 | 79 | 79 |
| CLK1 | 93 | 92 | 91 | 95 |
| CLK2 | 79 | 74 | 76 | 98 |
| CLK3 | 89 | 97 | 86 | 97 |
| CLK4 | 80 | 77 | 100 | 96 |
| CSF1R | 87 | 86 | 91 | 100 |
| CSF1R-autoinhibited | 75 | 85 | 72 | 87 |
| CSK | 85 | 100 | 70 | 97 |
| CSNK1A1 | 53 | 77 | 83 | 87 |
| CSNK1A1L | 93 | 100 | 97 | 100 |
| CSNK1D | 43 | 92 | 92 | 94 |
| CSNK1E | 53 | 100 | 57 | 98 |
| CSNK1G1 | 29 | 99 | 79 | 89 |
| CSNK1G2 | 7,1 | 80 | 91 | 100 |
| CSNK1G3 | 8 | 99 | 100 | 100 |
| CSNK2A1 | 90 | 62 | 89 | 81 |
| CSNK2A2 | 80 | 64 | 76 | 93 |
| CTK | 85 | 100 | 100 | 79 |
| DAPK1 | 90 | 100 | 93 | 93 |
| DAPK2 | 94 | 88 | 98 | 100 |
| DAPK3 | 100 | 86 | 96 | 95 |
| DCAMKL1 | 95 | 83 | 100 | 98 |
| DCAMKL2 | 92 | 87 | 96 | 100 |
| DCAMKL3 | 100 | 82 | 91 | 100 |
| DDR1 | 94 | 100 | 76 | 92 |
| DDR2 | 86 | 67 | 80 | 78 |
| DLK | 92 | 58 | 73 | 94 |
| DMPK | 52 | 100 | 94 | 83 |
| DMPK2 | 84 | 84 | 95 | 92 |
| DRAK1 | 97 | 100 | 97 | 94 |
| DRAK2 | 89 | 74 | 96 | 93 |
| DYRK1A | 0,1 | 96 | 98 | 100 |
| DYRK1B | 2,2 | 92 | 71 | 90 |
| DYRK2 | 49 | 50 | 69 | 67 |
| EGFR | 88 | 93 | 86 | 100 |
| EGFR(E746-A750del) | 71 | 80 | 47 | 97 |
| EGFR(G719C) | 92 | 84 | 100 | 100 |
| EGFR(G719S) | 100 | 82 | 100 | 100 |
| EGFR(L747-E749del, A750P) | 82 | 99 | 78 | 99 |
| EGFR(L747-S752del, P753S) | 80 | 59 | 85 | 97 |
| EGFR(L747-T751del,Sins) | 99 | 87 | 84 | 100 |
| EGFR(L858R) | 75 | 92 | 78 | 100 |
| EGFR(L858R,T790M) | 91 | 100 | 82 | 94 |

| | | | | |
|---|---|---|---|---|
| EGFR(L861Q) | 94 | 73 | 95 | 99 |
| EGFR(S752-I759del) | 78 | 86 | 95 | 99 |
| EGFR(T790M) | 91 | 100 | 89 | 99 |
| EIF2AK1 | 85 | 100 | 96 | 98 |
| EPHA1 | 85 | 100 | 79 | 94 |
| EPHA2 | 100 | 100 | 100 | 93 |
| EPHA3 | 86 | 95 | 100 | 100 |
| EPHA4 | 100 | 90 | 97 | 100 |
| EPHA5 | 100 | 86 | 100 | 100 |
| EPHA6 | 80 | 90 | 65 | 100 |
| EPHA7 | 92 | 83 | 92 | 91 |
| EPHA8 | 94 | 69 | 89 | 100 |
| EPHB1 | 100 | 83 | 98 | 82 |
| EPHB2 | 90 | 67 | 91 | 100 |
| EPHB3 | 79 | 92 | 80 | 100 |
| EPHB4 | 78 | 100 | 100 | 100 |
| EPHB6 | 100 | 53 | 90 | 100 |
| ERBB2 | 83 | 98 | 99 | 100 |
| ERBB3 | 100 | 80 | 100 | 88 |
| ERBB4 | 78 | 89 | 81 | 100 |
| ERK1 | 100 | 86 | 100 | 100 |
| ERK2 | 90 | 80 | 100 | 79 |
| ERK3 | 83 | 64 | 96 | 94 |
| ERK4 | 100 | 99 | 100 | 100 |
| ERK5 | 91 | 99 | 100 | 94 |
| ERK8 | 74 | 94 | 84 | 86 |
| ERN1 | 81 | 97 | 91 | 76 |
| FAK | 91 | 83 | 92 | 74 |
| FER | 100 | 92 | 98 | 100 |
| FES | 87 | 92 | 100 | 100 |
| FGFR1 | 89 | 93 | 73 | 98 |
| FGFR2 | 100 | 90 | 100 | 100 |
| FGFR3 | 95 | 80 | 100 | 100 |
| FGFR3(G697C) | 97 | 96 | 92 | 92 |
| FGFR4 | 100 | 93 | 100 | 100 |
| FGR | 74 | 100 | 63 | 91 |
| FLT1 | 96 | 100 | 100 | 87 |
| FLT3 | 76 | 64 | 81 | 94 |
| FLT3(D835H) | 96 | 78 | 76 | 93 |
| FLT3(D835V) | 100 | 91 | 100 | 100 |
| FLT3(D835Y) | 82 | 100 | 89 | 100 |
| FLT3(ITD) | 96 | 100 | 100 | 100 |
| FLT3(ITD,D835V) | 89 | 63 | 83 | 96 |
| FLT3(ITD,F691L) | 67 | 80 | 56 | 86 |
| FLT3(K663Q) | 93 | 100 | 92 | 95 |
| FLT3(N841I) | 97 | 44 | 96 | 100 |
| FLT3(R834Q) | 87 | 97 | 80 | 63 |
| FLT3-autoinhibited | 99 | 100 | 100 | 100 |
| FLT4 | 77 | 76 | 74 | 75 |
| FRK | 100 | 100 | 81 | 100 |
| FYN | 100 | 95 | 85 | 100 |
| GAK | 100 | 97 | 50 | 71 |
| GCN2(Kin.Dom.2,S808G) | 96 | 47 | 96 | 100 |
| GRK1 | 75 | 84 | 94 | 89 |
| GRK2 | 88 | 89 | 90 | 85 |
| GRK3 | 88 | 54 | 62 | 85 |
| GRK4 | 79 | 86 | 90 | 93 |
| GRK7 | 32 | 81 | 81 | 83 |
| GSK3A | 54 | 55 | 59 | 56 |
| GSK3B | 100 | 87 | 100 | 100 |
| HASPIN | 100 | 100 | 57 | 92 |
| HCK | 100 | 86 | 100 | 98 |
| HIPK1 | 75 | 73 | 68 | 85 |
| HIPK2 | 90 | 87 | 83 | 87 |

| | | | | |
|---|---|---|---|---|
| HIPK3 | 86 | 88 | 89 | 94 |
| HIPK4 | 83 | 86 | 80 | 62 |
| HPK1 | 85 | 92 | 82 | 99 |
| HUNK | 92 | 68 | 92 | 91 |
| ICK | 98 | 100 | 91 | 100 |
| IGF1R | 94 | 78 | 95 | 99 |
| IKK-alpha | 93 | 84 | 78 | 96 |
| IKK-beta | 100 | 88 | 93 | 100 |
| IKK-epsilon | 100 | 79 | 100 | 100 |
| INSR | 85 | 62 | 94 | 95 |
| INSRR | 85 | 62 | 96 | 99 |
| IRAK1 | 79 | 86 | 85 | 93 |
| IRAK3 | 87 | 91 | 93 | 96 |
| IRAK4 | 94 | 71 | 93 | 99 |
| ITK | 100 | 78 | 95 | 97 |
| JAK1(JH1domain-catalytic) | 88 | 79 | 97 | 100 |
| JAK1(JH2domain-pseudokinase) | 97 | 2,2 | 72 | 99 |
| JAK2(JH1domain-catalytic) | 97 | 66 | 85 | 88 |
| JAK3(JH1domain-catalytic) | 97 | 74 | 82 | 97 |
| JNK1 | 88 | 100 | 88 | 85 |
| JNK2 | 69 | 100 | 69 | 70 |
| JNK3 | 77 | 100 | 86 | 76 |
| KIT | 94 | 80 | 71 | 92 |
| KIT(A829P) | 85 | 57 | 69 | 91 |
| KIT(D816H) | 87 | 72 | 71 | 85 |
| KIT(D816V) | 84 | 100 | 69 | 100 |
| KIT(L576P) | 96 | 87 | 61 | 74 |
| KIT(V559D) | 100 | 85 | 67 | 87 |
| KIT(V559D,T670I) | 78 | 89 | 95 | 96 |
| KIT(V559D,V654A) | 93 | 100 | 88 | 91 |
| KIT-autoinhibited | 100 | 85 | 100 | 100 |
| LATS1 | 76 | 95 | 65 | 97 |
| LATS2 | 100 | 63 | 100 | 100 |
| LCK | 100 | 84 | 95 | 100 |
| LIMK1 | 99 | 93 | 100 | 93 |
| LIMK2 | 77 | 83 | 92 | 93 |
| LKB1 | 64 | 41 | 90 | 93 |
| LOK | 99 | 83 | 88 | 88 |
| LRRK2 | 91 | 73 | 88 | 100 |
| LRRK2(G2019S) | 95 | 86 | 93 | 79 |
| LTK | 100 | 76 | 100 | 95 |
| LYN | 97 | 75 | 83 | 100 |
| LZK | 77 | 88 | 83 | 74 |
| MAK | 75 | 76 | 93 | 82 |
| MAP3K1 | 88 | 86 | 95 | 93 |
| MAP3K15 | 78 | 74 | 100 | 93 |
| MAP3K2 | 100 | 73 | 97 | 90 |
| MAP3K3 | 99 | 79 | 68 | 100 |
| MAP3K4 | 97 | 95 | 100 | 95 |
| MAP4K2 | 70 | 91 | 84 | 100 |
| MAP4K3 | 91 | 69 | 100 | 100 |
| MAP4K4 | 93 | 100 | 100 | 97 |
| MAP4K5 | 98 | 100 | 100 | 100 |
| MAPKAPK2 | 67 | 99 | 94 | 94 |
| MAPKAPK5 | 86 | 77 | 89 | 97 |
| MARK1 | 97 | 100 | 100 | 96 |
| MARK2 | 75 | 88 | 100 | 100 |
| MARK3 | 34 | 86 | 60 | 46 |
| MARK4 | 80 | 100 | 82 | 91 |
| MAST1 | 99 | 56 | 97 | 100 |
| MEK1 | 100 | 82 | 97 | 100 |
| MEK2 | 90 | 67 | 76 | 83 |
| MEK3 | 96 | 61 | 86 | 92 |
| MEK4 | 92 | 88 | 88 | 100 |

| | | | | |
|---|---|---|---|---|
| MEK5 | 73 | 51 | 41 | 91 |
| MEK6 | 75 | 100 | 46 | 57 |
| MELK | 95 | 68 | 81 | 93 |
| MERTK | 100 | 84 | 100 | 100 |
| MET | 100 | 77 | 100 | 100 |
| MET(M1250T) | 100 | 97 | 100 | 96 |
| MET(Y1235D) | 83 | 100 | 86 | 100 |
| MINK | 66 | 84 | 79 | 89 |
| MKK7 | 81 | 87 | 73 | 77 |
| MKNK1 | 86 | 72 | 62 | 80 |
| MKNK2 | 75 | 100 | 74 | 81 |
| MLCK | 100 | 92 | 75 | 89 |
| MLK1 | 100 | 82 | 100 | 100 |
| MLK2 | 88 | 99 | 96 | 100 |
| MLK3 | 100 | 98 | 98 | 100 |
| MRCKA | 88 | 82 | 98 | 54 |
| MRCKB | 100 | 96 | 97 | 100 |
| MST1 | 64 | 73 | 80 | 62 |
| MST1R | 79 | 71 | 95 | 83 |
| MST2 | 100 | 0 | 84 | 94 |
| MST3 | 89 | 96 | 77 | 100 |
| MST4 | 100 | 99 | 100 | 100 |
| MTOR | 88 | 0 | 89 | 85 |
| MUSK | 76 | 95 | 96 | 92 |
| MYLK | 95 | 45 | 83 | 96 |
| MYLK2 | 92 | 100 | 11 | 88 |
| MYLK4 | 89 | 100 | 78 | 100 |
| MYO3A | 81 | 93 | 82 | 100 |
| MYO3B | 100 | 85 | 100 | 100 |
| NDR1 | 76 | 73 | 80 | 80 |
| NDR2 | 83 | 71 | 76 | 82 |
| NEK1 | 73 | 100 | 37 | 53 |
| NEK10 | 82 | 1,7 | 81 | 82 |
| NEK11 | 85 | 93 | 82 | 78 |
| NEK2 | 90 | 99 | 91 | 98 |
| NEK3 | 79 | 95 | 91 | 95 |
| NEK4 | 97 | 64 | 83 | 100 |
| NEK5 | 77 | 77 | 94 | 85 |
| NEK6 | 87 | 83 | 100 | 99 |
| NEK7 | 94 | 88 | 99 | 100 |
| NEK9 | 100 | 88 | 100 | 100 |
| NIK | 98 | 69 | 97 | 94 |
| NIM1 | 76 | 100 | 83 | 76 |
| NLK | 90 | 64 | 60 | 97 |
| OSR1 | 93 | 100 | 89 | 82 |
| p38-alpha | 89 | 98 | 92 | 100 |
| p38-beta | 99 | 96 | 93 | 100 |
| p38-delta | 93 | 92 | 97 | 93 |
| p38-gamma | 100 | 38 | 100 | 100 |
| PAK1 | 89 | 82 | 84 | 96 |
| PAK2 | 94 | 97 | 99 | 91 |
| PAK3 | 100 | 75 | 100 | 95 |
| PAK4 | 93 | 90 | 96 | 100 |
| PAK6 | 99 | 100 | 100 | 99 |
| PAK7 | 90 | 93 | 99 | 100 |
| PCTK1 | 88 | 100 | 84 | 89 |
| PCTK2 | 88 | 100 | 88 | 99 |
| PCTK3 | 98 | 84 | 92 | 100 |
| PDGFRA | 91 | 91 | 82 | 100 |
| PDGFRB | 94 | 83 | 57 | 96 |
| PDPK1 | 100 | 72 | 100 | 99 |
| PFCDPK1(P.falciparum) | 92 | 82 | 78 | 99 |
| PFPK5(P.falciparum) | 95 | 84 | 99 | 92 |
| PFTAIRE2 | 96 | 83 | 100 | 100 |

| | | | | |
|---|---|---|---|---|
| PFTK1 | 100 | 97 | 96 | 99 |
| PHKG1 | 97 | 94 | 97 | 100 |
| PHKG2 | 88 | 83 | 88 | 98 |
| PIK3C2B | 51 | 9,6 | 39 | 21 |
| PIK3C2G | 100 | 0 | 71 | 8 |
| PIK3CA | 100 | 0 | 95 | 8,2 |
| PIK3CA(C420R) | 74 | 0,05 | 56 | 2,9 |
| PIK3CA(E542K) | 81 | 0 | 82 | 6,2 |
| PIK3CA(E545A) | 88 | 0,65 | 64 | 3,4 |
| PIK3CA(E545K) | 81 | 0,15 | 79 | 6,4 |
| PIK3CA(H1047L) | 56 | 0 | 43 | 0 |
| PIK3CA(H1047Y) | 99 | 0 | 70 | 13 |
| PIK3CA(I800L) | 83 | 0,4 | 64 | 8,5 |
| PIK3CA(M1043I) | 94 | 0,85 | 83 | 6,7 |
| PIK3CA(Q546K) | 62 | 0 | 62 | 4,6 |
| PIK3CB | 99 | 0 | 12 | 12 |
| PIK3CD | 3,9 | 0 | 0,1 | 3,9 |
| PIK3CG | 21 | 0,05 | 15 | 2,5 |
| PIK4CB | 100 | 7,2 | 100 | 2,2 |
| PIKFYVE | 84 | 86 | 27 | 7,2 |
| PIM1 | 100 | 86 | 95 | 100 |
| PIM2 | 100 | 78 | 100 | 100 |
| PIM3 | 93 | 100 | 89 | 100 |
| PIP5K1A | 96 | 100 | 100 | 100 |
| PIP5K1C | 68 | 70 | 40 | 100 |
| PIP5K2B | 76 | 98 | 99 | 99 |
| PIP5K2C | 6,3 | 100 | 94 | 8,3 |
| PKAC-alpha | 94 | 91 | 100 | 100 |
| PKAC-beta | 100 | 93 | 100 | 99 |
| PKMYT1 | 99 | 100 | 99 | 100 |
| PKN1 | 88 | 77 | 91 | 100 |
| PKN2 | 100 | 100 | 100 | 100 |
| PKNB(M.tuberculosis) | 93 | 81 | 87 | 100 |
| PLK1 | 76 | 61 | 81 | 83 |
| PLK2 | 85 | 93 | 93 | 96 |
| PLK3 | 100 | 89 | 100 | 100 |
| PLK4 | 76 | 88 | 73 | 89 |
| PRKCD | 91 | 91 | 75 | 97 |
| PRKCE | 100 | 97 | 91 | 100 |
| PRKCH | 98 | 83 | 100 | 97 |
| PRKCI | 67 | 87 | 79 | 69 |
| PRKCQ | 96 | 90 | 89 | 96 |
| PRKD1 | 100 | 100 | 95 | 96 |
| PRKD2 | 85 | 78 | 97 | 94 |
| PRKD3 | 100 | 100 | 95 | 100 |
| PRKG1 | 93 | 95 | 100 | 87 |
| PRKG2 | 62 | 100 | 53 | 67 |
| PRKR | 94 | 100 | 100 | 100 |
| PRKX | 87 | 100 | 96 | 100 |
| PRP4 | 100 | 64 | 100 | 100 |
| PYK2 | 97 | 89 | 94 | 98 |
| QSK | 100 | 61 | 100 | 100 |
| RAF1 | 66 | 99 | 51 | 62 |
| RET | 79 | 89 | 79 | 98 |
| RET(M918T) | 79 | 72 | 66 | 71 |
| RET(V804L) | 72 | 76 | 74 | 71 |
| RET(V804M) | 92 | 98 | 96 | 92 |
| RIOK1 | 99 | 100 | 96 | 97 |
| RIOK2 | 96 | 30 | 99 | 100 |
| RIOK3 | 96 | 99 | 95 | 95 |
| RIPK1 | 79 | 78 | 88 | 92 |
| RIPK2 | 71 | 74 | 89 | 100 |
| RIPK4 | 80 | 100 | 78 | 80 |
| RIPK5 | 100 | 75 | 100 | 100 |

| | | | | |
|---|---|---|---|---|
| ROCK1 | 92 | 78 | 100 | 100 |
| ROCK2 | 74 | 77 | 71 | 80 |
| ROS1 | 93 | 97 | 90 | 100 |
| RPS6KA4(Kin.Dom.1-N-terminal) | 87 | 100 | 91 | 82 |
| RPS6KA4(Kin.Dom.2-C-terminal) | 91 | 58 | 84 | 96 |
| RPS6KA5(Kin.Dom.1-N-terminal) | 100 | 75 | 100 | 98 |
| RPS6KA5(Kin.Dom.2-C-terminal) | 94 | 83 | 98 | 90 |
| RSK1(Kin.Dom.1-N-terminal) | 80 | 97 | 100 | 100 |
| RSK1(Kin.Dom.2-C-terminal) | 88 | 100 | 86 | 92 |
| RSK2(Kin.Dom.1-N-terminal) | 100 | 84 | 81 | 100 |
| RSK2(Kin.Dom.2-C-terminal) | 92 | 76 | 100 | 99 |
| RSK3(Kin.Dom.1-N-terminal) | 94 | 76 | 91 | 100 |
| RSK3(Kin.Dom.2-C-terminal) | 76 | 96 | 91 | 96 |
| RSK4(Kin.Dom.1-N-terminal) | 100 | 93 | 100 | 100 |
| RSK4(Kin.Dom.2-C-terminal) | 83 | 94 | 73 | 97 |
| S6K1 | 100 | 94 | 99 | 96 |
| SBK1 | 98 | 100 | 100 | 100 |
| SGK | 100 | 69 | 99 | 96 |
| SgK110 | 80 | 100 | 76 | 100 |
| SGK2 | 90 | 61 | 100 | 92 |
| SGK3 | 89 | 100 | 100 | 100 |
| SIK | 93 | 85 | 66 | 95 |
| SIK2 | 88 | 74 | 85 | 100 |
| SLK | 100 | 100 | 78 | 100 |
| SNARK | 67 | 100 | 49 | 71 |
| SNRK | 100 | 100 | 98 | 99 |
| SRC | 100 | 80 | 100 | 100 |
| SRMS | 79 | 80 | 68 | 92 |
| SRPK1 | 100 | 82 | 100 | 100 |
| SRPK2 | 90 | 100 | 93 | 97 |
| SRPK3 | 100 | 84 | 100 | 100 |
| STK16 | 91 | 69 | 92 | 100 |
| STK33 | 99 | 66 | 97 | 83 |
| STK35 | 86 | 86 | 100 | 100 |
| STK36 | 95 | 69 | 91 | 100 |
| STK39 | 95 | 63 | 95 | 99 |
| SYK | 93 | 99 | 91 | 100 |
| TAK1 | 70 | 77 | 80 | 87 |
| TAOK1 | 40 | 91 | 65 | 73 |
| TAOK2 | 95 | 89 | 81 | 93 |
| TAOK3 | 76 | 72 | 88 | 99 |
| TBK1 | 85 | 77 | 57 | 78 |
| TEC | 90 | 81 | 92 | 98 |
| TESK1 | 95 | 76 | 73 | 96 |
| TGFBR1 | 93 | 99 | 63 | 88 |
| TGFBR2 | 89 | 96 | 91 | 100 |
| TIE1 | 81 | 71 | 90 | 89 |
| TIE2 | 91 | 93 | 75 | 95 |
| TLK1 | 94 | 80 | 83 | 88 |
| TLK2 | 92 | 100 | 82 | 96 |
| TNIK | 80 | 89 | 96 | 100 |
| TNK1 | 100 | 67 | 94 | 100 |
| TNK2 | 100 | 100 | 100 | 100 |
| TNNI3K | 76 | 84 | 95 | 100 |
| TRKA | 83 | 56 | 56 | 89 |
| TRKB | 76 | 90 | 72 | 76 |
| TRKC | 88 | 59 | 90 | 92 |
| TRPM6 | 100 | 72 | 87 | 100 |
| TSSK1B | 80 | 89 | 91 | 93 |
| TSSK3 | 89 | 82 | 89 | 96 |
| TTK | 89 | 95 | 88 | 85 |
| TXK | 100 | 83 | 100 | 100 |
| TYK2(JH1domain-catalytic) | 89 | 61 | 96 | 92 |
| TYK2(JH2domain-pseudokinase) | 90 | 100 | 32 | 85 |

| | | | | |
|---|---|---|---|---|
| TYRO3 | 100 | 72 | 100 | 100 |
| ULK1 | 94 | 100 | 90 | 81 |
| ULK2 | 96 | 86 | 92 | 88 |
| ULK3 | 95 | 92 | 94 | 76 |
| VEGFR2 | 93 | 73 | 83 | 100 |
| VPS34 | 87 | 0,95 | 31 | 77 |
| VRK2 | 76 | 48 | 79 | 96 |
| WEE1 | 86 | 94 | 100 | 99 |
| WEE2 | 96 | 100 | 95 | 94 |
| WNK1 | 83 | 98 | 92 | 100 |
| WNK2 | 99 | 83 | 95 | 96 |
| WNK3 | 100 | 100 | 100 | 100 |
| WNK4 | 71 | 97 | 60 | 77 |
| YANK1 | 94 | 74 | 71 | 100 |
| YANK2 | 98 | 92 | 87 | 90 |
| YANK3 | 81 | 96 | 63 | 89 |
| YES | 100 | 100 | 88 | 96 |
| YSK1 | 60 | 81 | 24 | 32 |
| YSK4 | 73 | 67 | 66 | 85 |
| ZAK | 100 | 92 | 100 | 100 |
| ZAP70 | 96 | 72 | 92 | 89 |
