## Supplementary Table 4 for "Functional testing of PI3K inhibitors stratifies responders to idelalisib and identifies treatment vulnerabilities in idelalisib-refractory/intolerant chronic lymphocytic leukemia"

**Supplementary Table 4.** Patient characteristics

| Patient ID | Gender | Age | Binet stage | IGVH | FISH/TP53 | Treatment at procurement | Treatment prior to procurement | Samples collected (if more than one) | Treatment status as indicated in figures |
|---|---|---|---|---|---|---|---|---|---|
| CLL001D | m | 68 | A | M | n.e. | None | None | | |
| CLL002D | m | 66 | A | UM | n.e. | None | None | | Treatment naive |
| CLL003D | m | 73 | A | M | n.e. | None | None | | Treatment naive |
| CLL103 | m | 62 | C | UM | del(13q14) | Idelalisib | FCR, idelalisib | | Idelalisib intolerant |
| CLL116 | m | 82 | A | M | n.e. | None | Radiation therapy for prostate cancer | | Treatment naive |
| CLL150 | f | 64 | C | UM | del(13q14), TP53 mutation | See Figure 5a | See Figure 5a | T0, T1, T2, T3, T4 | |
| CLL152 | f | 56 | B | UM | NOTCH1-mutation, no TP53-mutation | None | None | | |
| CLL153 | m | 51 | C | UM | del(11q22), trisomy 12, del(13q14) | None | None | | |
| CLL154 | m | 61 | A | M | n.e. | None | None | | |
| CLL159 | m | 66 | A | M | n.e. | None | None | | |
| CLL160 | m | 72 | A | M | 46,XY | None | None | | |
| CLL161 | m | 55 | A | M | n.e. | None | None | | |
| CLL163 | f | 66 | A | M | n.e. | None | None | | Treatment naive |
| CLL164 T1 | m | 73 | C | UM | del(13q14), TP53 mutation | None | Ibrutinib, idelalisib | | Idelalisib intolerant |
| CLL183 | m | 68 | C | M | Normal | None | FC, venetoclax, idelalisib | | Idelalisib intolerant |
| CLL185 | f | 58 | A | M | n.e. | None | None | | |
| CLL187 | m | 66 | A | M | n.e. | None | None | | |
| CLL200 | m | 64 | B | UM | del(13q14), del(17p) TP53-mutation, SF3B1-mutation and NRA-mutation | None | Idelalisib | | Idelalisib intolerant |
| CLL206 | f | 66 | C | UM | 46,XX, heterozygous del(13)(q14) | None | None | | Treatment naive |
| CLL211 | m | 80 | B | UM | trisomy 12, del(11q22) | None | BR, Idelalisib | | Idelalisib refractory |
| CLL216 | m | 58 | C | M | del(13)(q14) | None | C, radiation, FCR | | |
| CLL248 | m | 73 | B | M | del(13q14) + translocation involving 14q32 | None | None | | Treatment naive |
| CLL249 | f | 80 | A | n.e. | n.e. | None | None | | Treatment naive |
| CLL251 | f | 63 | C | M | Trisomy 12; NO TP53-mutation, no SF3B1-mutation, no NOTCH1-mutation | None | None | | |
| UMB2 HPA-0205-screening | m | 82 | n.e. | UM | Failed | None | BR, Cytoxan, ibrutinib, idelalisib | | Idelalisib intolerant |
| UMB3 HPA- | m | 72 | n.e. | M | del(11q22), TP53 mutation | None | FR, FCR, BR, ibrutinib, idelalisib + R, obinutuzumab | | Idelalisib intolerant |

| | | | | | | | | | |
|---|---|---|---|---|---|---|---|---|---|
| 0206-screening | | | | | | | | | |
| UMB5 NYA-0204-C2D1 | f | 78 | n.e. | M | Normal | Umbralisib | Idelalisib | | Idelalisib intolerant |
| UMB16 FMB-0202-screening | m | 80 | n.e. | M | Normal | None | Chl, R (2), CVP, BR, idelalisib | | Idelalisib intolerant |
| JB-0058 | m | 58, 61 | A, C | UM | del(11q22), del(13q14) | Idelalisib | FCR | Responding | Short-term responder |
| JB-0157 | f | 71, 71 | C | M | del(13q14), del(17p), TP53 mutation | Idelalisib (with OFA at initiation) | BR | Baseline, responding | Short-term responder |
| JB-0158 | m | 53, 55 | C, A | M | Normal | Idelalisib (with OFA at initiation) | BR | Baseline, responding | Short-term responder |
| JB-0197 | m | 80, 82 | A, C | UM | trisomy 12, del(17p) | Idelalisib | BR | Baseline, responding | Short-term responder |
| JB-0237 | m | 66, 66 | C, A | UM | del(13q14) | Idelalisib | None | Baseline, responding | Long-term responder |
| JB-0238 | m | 71, 72 | C, A | M | del(13q14) | Idelalisib | None | Baseline, responding | Long-term responder |
| JB-0244 | m | 80, 80 | C, A | UM | del(11q22), del(13q14), del(17p) | Idelalisib | None | Baseline, responding | Long-term responder |
