## Supplementary figures and images for "Functional testing of PI3K inhibitors stratifies responders to idelalisib and identifies treatment vulnerabilities in idelalisib-refractory/intolerant chronic lymphocytic leukemia"

### Supplementary Figures

#### Supplementary Figure 1

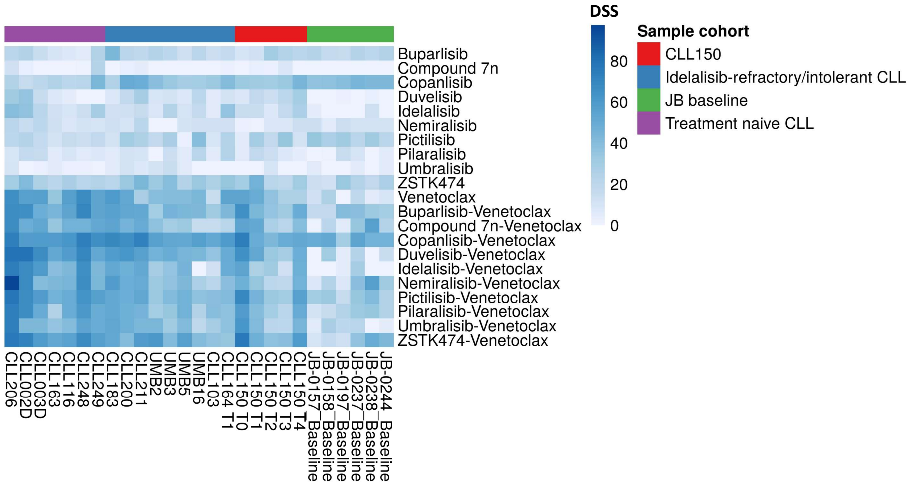

#### Supplementary Figure 2

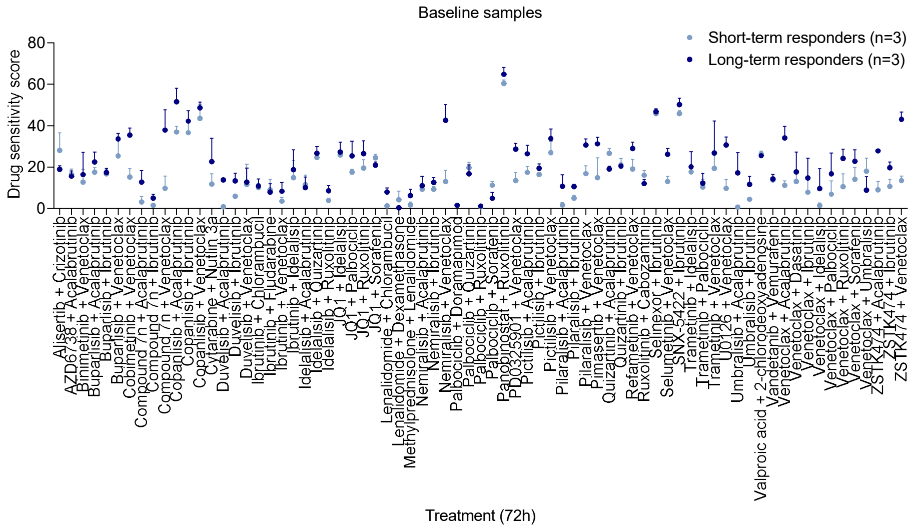
