## Supplementary Table 1 for "Functional testing of PI3K inhibitors stratifies responders to idelalisib and identifies treatment vulnerabilities in idelalisib-refractory/intolerant chronic lymphocytic leukemia"

**Supplementary Table 1.** Compound library

| Compound | Target | Supplier | Cat no | Solvent |
|---|---|---|---|---|
| 2-chlorodeoxyadenosine | Adenosine deaminase | AK Scientific | E997 | DMSO |
| Acalabrutinib | BTK | Selleck Chemicals LLC | S8116 | DMSO |
| Alisertib | Aurora A | Selleck Chemicals LLC | S1133 | DMSO |
| AZD6738 | ATR | TargetMol | T3338 | DMSO |
| Binimetinib | MEK1/2 | Selleck Chemicals LLC | S7007 | DMSO |
| Buparlisib | PI3K | Selleck Chemicals LLC | S2247 | DMSO |
| Cabozantinib | VEGFR2 | LC Laboratories | C-8901 | DMSO |
| Chlorambucil | Alkylating agent | Pure Chemistry Scientific | 815818 | DMSO |
| Cobimetinib | MEK1/2 | Selleck Chemicals LLC | S8041 | DMSO |
| Compound 7n | PI3K | Selleck Chemicals LLC | S8693 | DMSO |
| Copanlisib | PI3K | Selleck Chemicals LLC | S2802 | 5% TFA |
| Crizotinib | c-Met, ALK | LC Laboratories | C-7900 | DMSO |
| Cytarabine | DNA synthesis | Selleck Chemicals LLC | S1648 | H2O |
| Dasatinib | Abl, Src, c-Kit | LC Laboratories | D-3307 | DMSO |
| Dexamethasone | Interleukin receptor | Selleck Chemicals LLC | S1322 | DMSO |
| Doramapimod | p38 MAPK | LC Laboratories | D-2744 | DMSO |
| Duvelisib | PI3K | Selleck Chemicals LLC | S7028 | DMSO |
| Fludarabine | DNA synthesis, STAT1 | Selleck Chemicals LLC | S1491 | DMSO |
| Ibrutinib | BTK | LC Laboratories | S2680 | DMSO |
| Idelalisib | PI3K | LC Laboratories | S2226 | DMSO |
| JQ1 | BET bromodomain | Selleck Chemicals LLC | S7110 | DMSO |
| Lenalidomide | Immunomodulation | Selleck Chemicals LLC | S1029 | DMSO |
| Methylprednisolone | Glucocorticoid receptor | Selleck Chemicals LLC | S1733 | DMSO |
| Nemiralisib | PI3K | Selleck Chemicals LLC | S7937 | DMSO |
| Nutlin 3a | p53/MDM2 | Selleck Chemicals LLC | S8059 | DMSO |
| Palbociclib | CDK4/6 | LC Laboratories | P-7744 | DMSO |
| Panobinostat | HDAC | LC Laboratories | P-3703 | DMSO |
| PD0325901 | MEK1/2 | Selleck Chemicals LLC | S1036 | DMSO |
| Pictilisib | PI3K | Selleck Chemicals LLC | S1065 | DMSO |
| Pilaralisib | PI3K | Selleck Chemicals LLC | S7645 | DMSO |
| Pimasertib | MEK1/2 | Selleck Chemicals LLC | S1475 | DMSO |
| Quizartinib | FLT3 | LC Laboratories | Q-4747 | DMSO |
| Refametinib | MEK1/2 | Selleck Chemicals LLC | S1089 | DMSO |
| Ruxolitinib | JAK1/2 | LC Laboratories | R-6600 | DMSO |
| Selinexor | CRM1 | Cayman Europe | 18127.0 | DMSO |
| Selumetinib | MEK1/2 | Selleck Chemicals LLC | S1008 | DMSO |
| SNX-5422 | HSP90 | Cayman Europe | 18270.0 | DMSO |
| Sorafenib | Raf-1, B-Raf and VEGFR-2 | LC Laboratories | S-8599 | DMSO |
| Trametinib | MEK1/2 | LC Laboratories | T-8123 | DMSO |
| U0126 | MEK1/2 | Selleck Chemicals LLC | S1102 | DMSO |
| Umbralisib | PI3K | MedChemTronica | HY-12279 | DMSO |
| Valproic acid | Histone deacetylase | ENAMINE Ltd. | Z1511532065 | DMSO |
| Vandetanib | VEGFR2/3, EGFR | LC Laboratories | V-9402 | DMSO |
| Vemurafenib | B-Raf$^{V600E}$ | LC Laboratories | V-2800 | DMSO |
| Venetoclax | Bcl-2 | Selleck Chemicals LLC | S8048-5MG | DMSO |
| ZSTK474 | PI3K | Selleck Chemicals | S1072 | DMSO |
